# AXL inhibition reprograms tumor-associated macrophages to restore immune checkpoint blockade efficacy in a polarization-context-dependent manner

**DOI:** 10.1101/2025.07.30.667666

**Authors:** Amanda Kirane, David Lee, Saurabh Sharma, Elena Safrygina, Christopher J Applebee, Mamatha Serasanambati, Prabhjeet Singh, Jay Chadokiya, Emma Wagner, Alexander Meerlev, Daniel Delitto, Julian Padget, Banafshé Larijani, Emanual Maverakis

## Abstract

**Background:** Immune checkpoint blockade (ICB) achieves durable responses in approximately half of patients with advanced melanoma, but the mechanistic basis for resistance in the remaining patients remains incompletely defined. AXL tyrosine kinase has emerged as a candidate resistance mediator, yet clinical development of AXL inhibitors has yielded heterogeneous results in unselected patient populations, suggesting that cell-type and tumor microenvironment context may critically govern therapeutic response.

**Methods:** We characterized AXL expression across cell types in ICB-resistant melanoma using publicly available single-cell RNA sequencing datasets and The Cancer Genome Atlas (TCGA). In vivo efficacy was assessed in the YUMM1.7 PD-1-resistant syngeneic melanoma model using three pharmacologically distinct AXL inhibitors (warfarin, bemcentinib, and cabozantinib) as monotherapy and in combination with anti-PD-1 therapy. Tumor-associated macrophage (TAM) context-dependency was established using anti-CSF1R and anti-F4/80 depletion strategies. Functional PD-L1:PD-1 checkpoint interactions were quantified in tumor sections by immune Förster Resonance Energy Transfer (iFRET). TAM secretome reprogramming was characterized by 40-plex Luminex immunoassay in polarized RAW264.7 macrophages.

**Results:** AXL expression in ICB-resistant melanoma was predominantly localized to TAMs rather than tumor cells by single-cell analysis, with AXL+ TAMs distributed across both M1-like and M2-like phenotypic compartments. AXL inhibition significantly reduced tumor burden and synergized with anti-PD-1 therapy in vivo; however, efficacy was abolished by depletion of the monocyte-derived myeloid compartment (anti-CSF1R) and enhanced by depletion of tissue-resident TAMs (anti-F4/80), establishing TAM-context dependency. iFRET revealed a paradoxical gain of PD-L1:PD-1 interaction efficiency in anti-PD-1-treated resistant tumors, a functional resistance signature detectable by iFRET but not by PD-L1 expression that was reversed by AXL combination therapy. In vitro secretome profiling demonstrated that combination therapy reprograms macrophage secretome in a polarization-context-dependent manner, amplifying pro-inflammatory cytokines including IL-6 and GM-CSF in M2-like macrophages while selectively dampening T cell chemokine production.

**Conclusions:** These findings establish AXL as a TAM-resident immune target in ICB-resistant melanoma whose therapeutic relevance is governed by tumor-immune micronenvironment (TiME) macrophage composition rather than tumor cell AXL expression. TAM polarization contexture represents a candidate stratification axis for AXL inhibitor-based combination strategies, and iFRET-measured checkpoint interaction offers a functional complement to PD-L1 expression for monitoring resistance and response.

**Key Messages:** *What is already known on this topic:*

- AXL tyrosine kinase has been studied predominantly as a tumor-intrinsic mesenchymal marker and therapeutic target in melanoma, with prior clinical development of AXL inhibitors predicated on tumor cell AXL expression as the primary biomarker; the contribution of TAM-expressed AXL to ICB resistance and the dependence of therapeutic response on TiME composition have not been defined.

*What this study adds:*

- AXL expression in ICB-resistant melanoma is predominantly localized to tumor-associated macrophages distributed across both M1-like and M2-like phenotypic compartments, reframing AXL as a TAM immune target whose therapeutic relevance is governed by TiME macrophage composition rather than tumor cell AXL expression.
- Combination AXL inhibition and anti-PD-1 reprograms the macrophage secretome in a polarization-context-dependent manner and reverses a paradoxical gain of functional PD-L1:PD-1 checkpoint interaction in resistant tumors — a resistance signature detectable by spatial iFRET but not by PD-L1 expression.

*How this study might affect research, practice or policy:*

- The context-dependent effects of AXL inhibition (immunostimulatory in M2-heavy TiMEs, potentially counterproductive in M1-heavy TiMEs) indicate that TAM polarization profiling should be incorporated into clinical trial design for AXL inhibitor-based combinations, and that unselected enrollment may obscure meaningful efficacy signals in the subset most likely to benefit.
- iFRET-based quantification of functional checkpoint interaction represents a candidate dynamic biomarker for ICB resistance monitoring that is orthogonal to PD-L1 immunohistochemistry and may identify resistant patients who retain immunosuppressive checkpoint interactions that can be disrupted by AXL targeting.

**Graphical Abstract:** 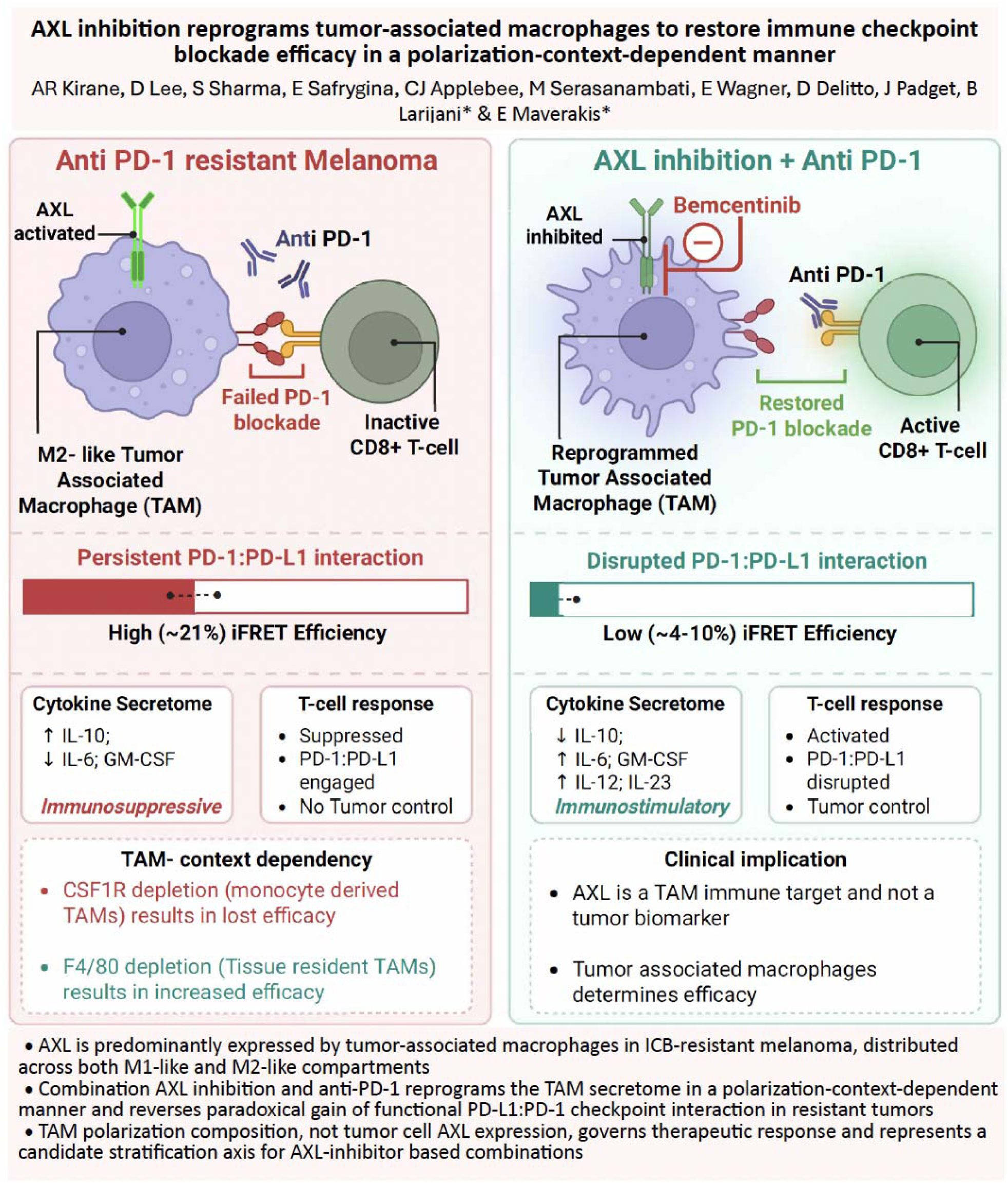

## Introduction

Immune checkpoint blockade (ICB) has revolutionized cancer care, proving highly effective in patients with advanced melanoma; however, only half of patients with melanoma show a meaningful response^1–3^. Melanoma presents a particular challenge due to its wide intra-and inter-tumoral heterogeneity that drives resistance to conventional therapies^4,5^. The receptor tyrosine kinase AXL, is a Tyro3/AXL/MerTK-family kinase that signals primarily through growth arrest-specific factor 6 (gas6)-ligand activated dimerization. When activated, AXL acts on various downstream pathways through PI3K/AKT, MAPK, and STAT3, leading to cell proliferation, survival, and immune modulation^6–9^. While AXL plays a vital physiologic role essential to cellular maintenance, it is expressed in multiple types of cancers on both tumor and non-tumor cells in the tumor immune microenvironment (TiME), such as dendritic cells, macrophages, fibroblasts, and NK cells^10^.

Our prior research has provided strong evidence that AXL blockade with either ligand inhibition via warfarin, a vitamin K carboxylase inhibitor, or the small molecule inhibitor bemcentinib (BGB324, Bem), can suppress tumor progression, metastasis, and drug resistance in some cancers^11,12^. We have recently shown that warfarin usage in patients with invasive melanoma is associated with improved melanoma-specific survival and overall survival^13^. High AXL expression by melanoma cells was also reported as a significant factor amongst a panel of mensenchymal-like tumor markers associated with ICB-resistance in stage IV melanoma^14–15^. AXL inhibition sensitize tumors to chemotherapies in breast and pancreatic murine models and to PD-1 checkpoint blockade in aPD-1-refractory leukemias^11,16^. Despite some clinical success in targeting AXL in multiple tumor types, AXL-related activity remains under preliminary exploration in melanoma^17^. Previously, a randomized phase Ib/II study of bemcentinib combined with pembrolizumab or dabrafenib/trametinib in metastatic melanoma (BGBIL006) did not demonstrate significant improvement in ORR, PFS, or OS in the unselected efficacy population, nor in pre-planned biomarker-defined subgroups based on tumor cell AXL expression^18–19^. Instead, high AXL expression in inflammatory cells, rather than tumor cells, emerged as a predictor of pembrolizumab response in exploratory analyses, a finding consistent with the hypothesis that TAM-compartment AXL expression, rather than tumor-intrinsic AXL, governs ICB sensitivity in melanoma.

Tumor-associated macrophages (TAMs) make up to 50% of tumor mass in melanoma and can exhibit a spectrum of phenotypes, ranging from immunostimulatory (M1-like) to immunosupproesive (M2-like)^20–21^. A high proportion of M2-like TAMs within the TiME contributes to tumor evasion of immune surveillance and is associated with a worse prognosis^12, 22–24^. The gas6/AXL axis is thought to classically drive an M2-like macrophage phenotype in a variety of systems and disease settings^25–26^. However, gas6/AXL signaling also amplifies an M1-like macrophage immunostimulatory response, and the overall effect of this equilibrium is dependent on the specific system or microenvironment^27^. Single-cell sequencing has demonstrated that M1/M2-like macrophage classification is highly limited with markers expressed in transitional states with highly plastic functions, making functional definition of AXL in TAM behaviors complex^28,-29^. In this study, we hypothesized that AXL inhibition can sensitize ICB-resistant melanoma to checkpoint therapy in the immunosuppressive TiME by reprogramming TAM functions and that AXL-driven responses are TiME-context dependent.

## Results

### The clinical landscape of AXL in melanoma

While AXL has been implicated in late-stage melanoma^15^, AXL expression is poorly across the clinical spectrum of melanoma. We first examined the association of AXL with melanoma survival by using the Cancer Genome Atlas (TCGA)-skin cutaneous melanoma (SKCM) dataset. We detected a significant relationship between increased AXL expression and increased overall survival (**Fig. 1A**) with no significant correlations observed with ligand gas6 or AXL-family receptor kinases Tyro3 and MerTK (**Supplementary Fig. S1A**). AXL expression trended in line with inflammatory infiltrate, such as CD68, CD8^+^ T cells, and Programmed Death Ligand-1 (PD-L1) expression(**Supplementary Fig. S1B**). Serum samples of untreated patients at our institution were screened for circulating levels of soluble AXL (sAXL). sAXL significantly increased in advanced disease stages, and was highest in patients with Stage IV disease (**Supplementary Fig. S1C**). Given the somewhat paradoxical finding that AXL expression correlated with survival in untreated melanoma tumors (**Fig. 1A)** but increased in advanced or PD-1 resistant melanoma, we aimed to resolve the expression of AXL by cell type using single cell sequencing datasets^14, 15, 29^.

**Figure 1.**
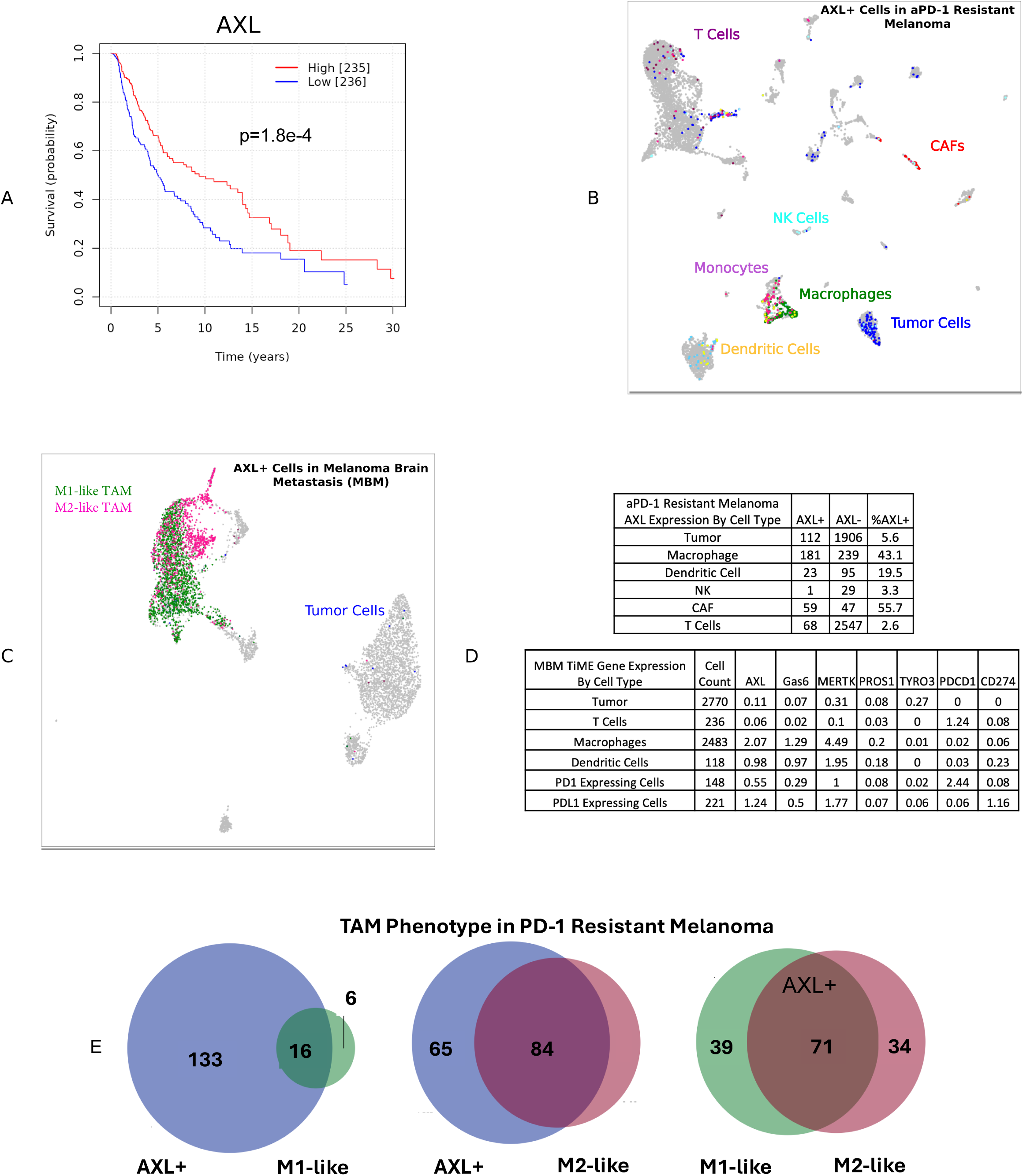
TAMs are the dominant source of AXL in melanoma. A) Kaplan-Meier survival curves created from TCGA-SKCM data demonstrate significant association of high AXL expression with increased overall survival in a broad cohort of melanoma. B) Single cell sequencing of aPD-1 refractory melanoma and C) melanoma brain metastasis (MBM), Uniform Manifold Approximation and Projection (UMAP) of AXL (expression >1) by cell type. D) Quantification of AXL+ cell types in aPD-1 resistant melanoma demonstrates predominant expression of AXL in tumors by tumor-associated macrophages (TAMs) with <10% of tumor cells demonstrating significant AXL expression. MBM demonstrate high AXL expression in TAMs with co-expression of PD-L1. E) AXL+ TAMs span both M1-like (CD86+/CD80+) and M2-like (CD163+/CD206+) TAMs with significant overlap in phenotypic markers by AXL+ TAMs.

Available single cell RNAseq datasets (GSE115978, GSM6022254) were examined for cell type-specific AXL expression, with focus on therapeutically challenging tumor types (e.g. melanoma brain metastases (MBM) and ICB-resistant melanoma) (**Fig. 1B,D**)^14,29^. Despite positive findings amongst tumor cells as previously reported by Jerby et al (GSE 115978), AXL was only expressed in a small subset of tumor cells. The predominant AXL expression was found in TAMs, in addition to smaller populations of TiME cells, such as dendritic cells, natural killer cells, and cancer-associated fibroblasts. Interestingly, we observed that AXL was strongly expressed in a subset of CD4^+^ T cells, which has not been previously reported but was consistently present in all examined datasets (**Fig. 1C,E**). Among TAMs, AXL expression was found to be equally present in M1-like (CD80/86+, TNF^+^, MHC class II) and M2-like (CD206^+^/MRC1^+^, CD163^+^, IL-10^+^, VEGF^+^) TAMs. Importantly, the most abundant source of PD-L1 in melanoma tumors was by TAMs, as opposed to tumor cells. AXL expression was significantly correlated to PD-L1 expression across cell types (**Fig. 1E**).

### AXL expression and function in melanoma cell lines

Melanoma models known to be ICB (aPD-1)-resistant (Yumm1.7, B16F10, B16F10-LN6 (LN6) mouse melanoma lines), and the ICB-sensitive (Yummer1.7)^30–31^, were screened for AXL expression by western blot and tested *in vitro* for AXL functional activity. Yumm1.7, B16F10, and LN6 demonstrated baseline AXL expression and p-AXL activation, while Yummer1.7 lines expressed minimal AXL (**Supplementary Fig. S2A**) and p-AXL levels were diminished by warfarin treatment (**Supplementary Fig. S2B**). As AXL has been associated with mesenchymal transition and invasive capacity in melanoma and other cancer types^12,32^, the impact of AXL inhibition on tumor cell behavior *in vitro* was measured by migration assay. Migration was significantly increased with gas6 stimulation and decreased with warfarin, both at baseline and in the setting of gas6 stimulation (**Supplementary Fig. S2C**). YUMM1.7 was used as the primary in vivo efficacy model for subsequent experiments.

### AXL inhibition augments aPD-1 efficacy in models of ICB-resistant melanoma

Orthotopic C57/Bl6 mouse models were treated with multiple strategies of AXL inhibition (warfarin, bemcentinib) as single agents or in combination with aPD-1 therapy (**Fig. 2A**). Limited data has suggested that AXL activity is associated with invasive, advanced melanoma but less present during proliferative tumor phases^30^. As such, we examined the impact of AXL inhibition prior to tumor implantation versus after tumor establishment, a factor proving critical in prior non-melanoma tumor models^12^. Here, however, no differences were observed with warfarin administration before or after tumor inoculation, but both warfarin-treated groups demonstrated significantly reduced tumor size compared to control (**Sup Fig. S2D-E**). Yumm1.7 tumors were not significantly controlled by aPD-1 therapy, consistent with prior reports^24,33^. Warfarin monotherapy significantly reduced tumor size compared to control (p<0.05) (**Fig. 2B**), while combination therapy with warfarin and aPD-1 yielded the smallest tumors (p<0.05) (**Fig. 2B**). Studies were repeated with Bem, as monotherapy or in combination with aPD-1, and again demonstrated significantly reduced tumor size compared to control and aPD-1 alone (**Fig. 2C**). Similar tumor control was observed with cabozantinib, a multi-kinase inhibitor with AXL inhibitory activity, consistent with AXL pathway engagement as the relevant mechanism (**Supplementary Fig. S2F**).

**Figure 2.**
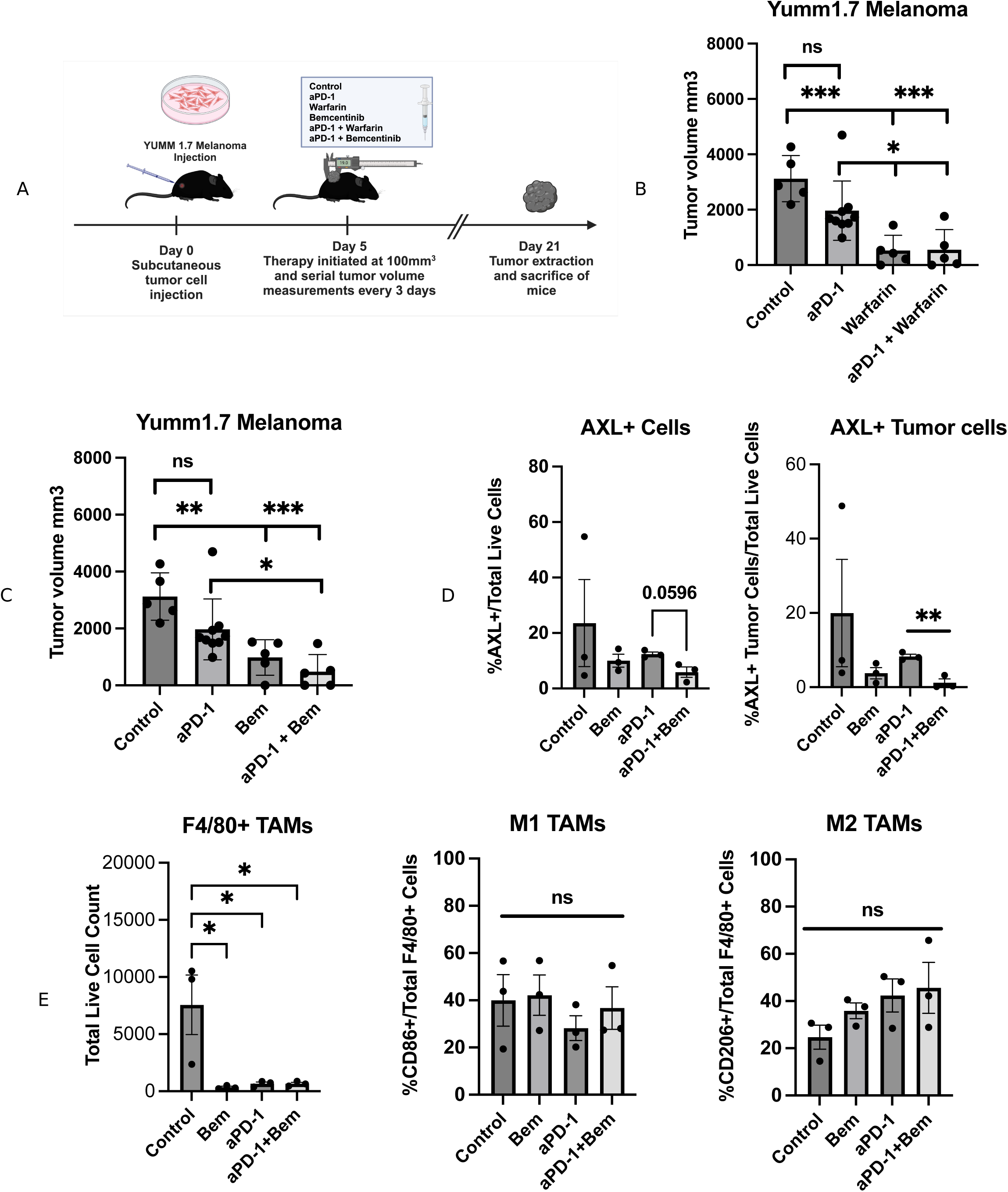
AXL inhibition synergizes with aPD-1 therapy. A) Schematic of mouse model of aPD-1-resistant melanoma and therapeutic groups, n=5/group. warfarin and bemcentinib (Bem) treatment groups were performed in triplicate. B,C) Warfarin and bemcentinib were effective as single agent therapy and significantly reduced tumor size in combination with aPD-1 therapy, compared to aPD-1 therapy alone (p<0.05). D) Combination therapy reduced the AXL+ tumor cell population (p<0.01) but overall reduction AXL+ population did not reach significance (p=0.0596). E) Global F4/80+ TAM depletion was observed in all treatment groups (p<0.05) but with no significant changes in TAM-phenotypic expression markers.

### TiME Shifts with AXL and PD-1 Inhibition

Tumors were further examined by flow cytometry for T cell populations (CD4, CD8), AXL expression, and myeloid constitution (F480^+^ TAMs, M1-like (CD86+) and M2-like (CD206+) phenotypes, and AXL^+^ cells). AXL^+^ cells represented an overall small population of tumor cells. Decrease in the AXL^+^ population was observed with combination aPD-1 and Bem compared to those treated with aPD-1 monotherapy, however differences did not reach significance (p=0.0596) (**Fig 2D**). F4/80^+^ TAMs were overall reduced in all treatment groups; however, our group observed that, unlike prior reports of AXL inhibition in pancreatic cancers^11^, neither TAM phenotype (M1-like or M2-like) nor AXL+TAMs differed significantly between groups (**Fig 2E, Supplementary Fig. S3A**). T cell population (total CD3+ T cell, CD4+ and CD8+ subpopulations) differences were also insignificant (**Supplementary Fig. S3B**).

### Functional checkpoint interactions demonstrate restoration of PD-L1/PD-1 blockade with AXL inhibition

Given that AXL inhibition significantly sensitized YUMM1.7 tumors to aPD-1 therapy without significant changes in T cell populations, TAM-phenotypic expression markers, or AXL expression, we examined functional PD-L1:PD-1 checkpoint engagement using immune Förster Resonance Energy Transfer (iFRET). iFRET quantifies receptor-ligand interaction at 1-10 nm, the actual interaction distance, offering substantially improved resolution over PD-L1 immunohistochemistry, which operates at approximately 70 nm and detects protein presence rather than molecular engagement (**Fig. 4A**). We have previously shown that PD-L1 expression does not correlate with PD-L1:PD-1 interaction efficiency (E*f*), and that ICB-resistant tumors gain E*f* while responding tumors show loss of E*f* in patient samples^34–36^.

iFRET measured across viable tumor sections revealed a striking and paradoxical pattern. Control tumors showed low baseline E*f*, consistent with minimal functional checkpoint engagement. Warfarin monotherapy produced intermediate E*f* of approximately 10%. aPD-1-treated tumors showed the highest E*f* of any group, with a median of 21%, significantly above control (p<0.001), indicating that the antibody failed to disrupt checkpoint engagement in this resistant model (**Fig. 4B**). Combination therapy restored E*f* to baseline range (**Fig. 4B**). PD-L1 expression did not change significantly across treatment groups, with the highest median expression in the combination arm, further demonstrating that expression and engagement are independent variables in this model (**Fig. 4C**).

This gain-of-interaction pattern recapitulates the functional resistance signature previously identified in ICB-treated melanoma patients, where non-responding patients gained E*f* and responding patients showed loss of E*f*^33^. iFRET measures PD-L1:PD-1 interactions across within tumor sections without cell-type resolution. Based on single-cell sequencing identifying TAMs as the dominant PD-L1-expressing population in these tumors, TAM-T cell interfaces are the most probable primary contributor to the observed interaction gain. iFRET co-registration with multiplex immunofluorescence^35^ is ongoing in patient-derived melanoma tissues to test this directly.

### Selective Depletion of TAMs Determines AXL Inhibitor Efficacy

We employed two models of TAM-depletion to determine the impact of AXL-targeting in the TiME on overall response to therapies. Anti-CSF1R or anti-F4/80 antibodies were administered prior to tumor inoculation to selectively deplete monocyte-derived or tissue-resident TAM compartments respectively (**Fig. 3A**). CSF1R depletion abolished differences in tumor size across treatment groups (**Fig. 3B**) and produced significantly accelerated tumor growth requiring early sacrifice at Day 14 (**Fig. 3C**). F4/80 depletion did not affect control or aPD-1 monotherapy tumor sizes, but sensitized tumors to Bem (p<0.01) and produced the greatest tumor reduction with combination aPD-1+Bem vs. both F4/80-depleted controls and non-depleted combination animals (p<0.05) (**Fig. 3D**).

**Figure 3.**
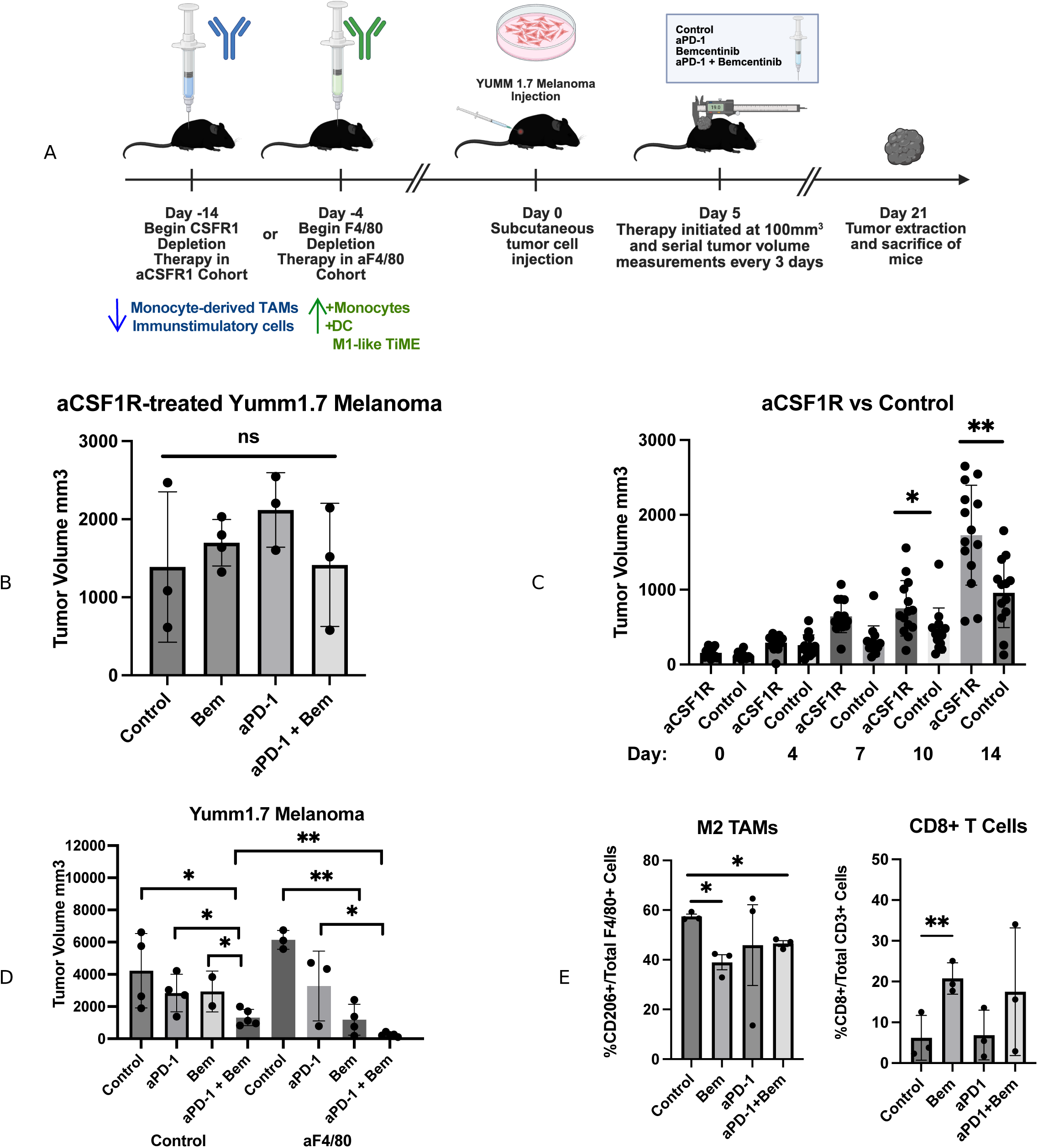
AXL and PD-1 efficacy in vivo is context-dependent. A) Schematic of macrophage depletion approaches. Anti-CSF1R antibody administration began 2 weeks before tumor injection, anti-F4/80 antibodies were administered 3 days prior to tumor injection and maintained throughout the duration of therapy. B) Final tumor volumes were not significantly different among therapy groups in CSF1R depleted mice. C) Accelerated tumor growth was observed in anti-CSF1R treated animals with significant size differences observed among control animals at day 10 and persisting at day of sacrifice (p<0.05). D) F4/80 depletion, alternatively, sensitized tumors to single agent bemcentinib (p<0.01) but not aPD-1 therapy. Combination therapy significantly reduced tumor size vs. aPD-1 monotherapy (p<0.05) and vs. non-depleted controls (p<0.001). E) In F4/80 depleted animals, there was a significant reduction in M2 TAMs (CD206+) in bemcentinib-treated compared with control (p<0.05) and bemcentinib increased CD8+ T cell infiltrate (p<0.01 vs. control). Anti-CSF1R depletes the monocyte-derived myeloid compartment, removing the primary immunostimulatory population available for reprogramming. Anti-F4/80 depletion removes tissue-resident M2-like TAMs while preserving monocytes, dendritic cells, and other pro-inflammatory innate populations, leaving the TiME relatively M1-intact.

**Figure 4.**
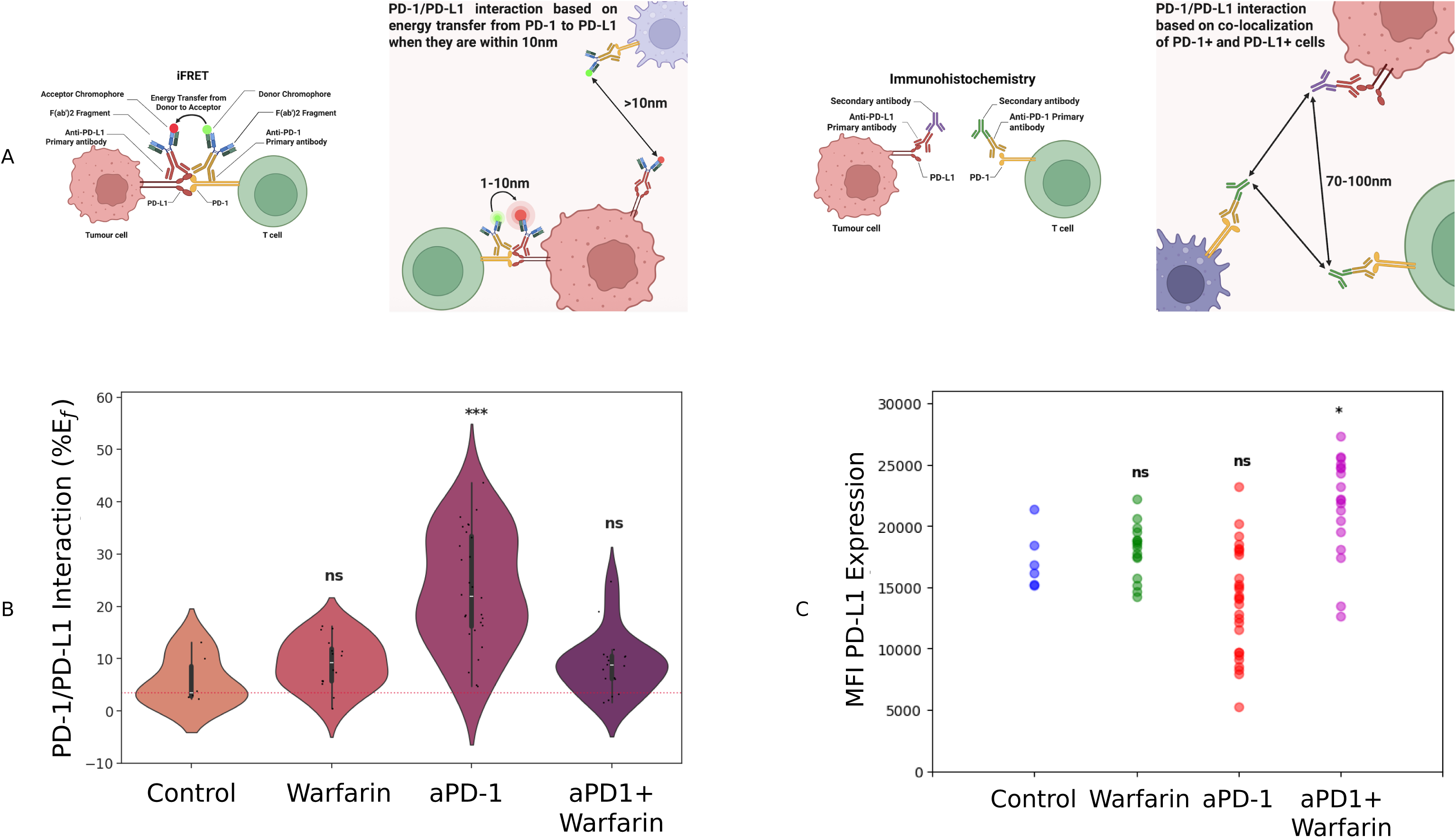
AXL inhibition disrupts PD-L1:PD-1 checkpoint interaction to restore checkpoint blockade. A) Schematic of iFRET assay quantifying PD-1/PD-L1 checkpoint interaction at the functional interaction distance (<10nm) compared to immunohistochemistry (70nm) and proximity ligation assay (40nm, not shown). Representative high and low interaction examples shown. B) iFRET-measured PD-1/PD-L1 checkpoint interactions in YUMM1.7 melanoma tumors, displayed as violin plot, demonstrate decrease in PD-1/PD-L1 interaction in the setting of warfarin therapy with highest interaction observed in aPD-1 resistant tumors (p<0.001). C) PD-L1 expression does not correlate to PD-1/PD-L1 interaction with high PD-L1 expression measured in combination treatment group (p=0.018). iFRET measurements (E*f* values) reflect aggregate PD-L1:PD-1 interaction signal across the viable tumor section.

### TiME Impact of Myeloid Cell Depletion

Tumors were analyzed with flow cytometry to investigate shifts in both macrophage and T cell populations. Overall, TAMs were depleted in the anti-CSF1R and anti-F4/80 groups, correlating to a decrease in AXL+ cells (**Supplementary Fig. S4A**). Anti-CSF1R depletion broadly targets the monocyte-derived myeloid compartment, removing the primary immunostimulatory macrophage population available for reprogramming. Anti-F4/80 depletion selectively removes tissue-resident macrophages, predominantly M2-like in melanoma, while preserving monocytes, dendritic cells, and other pro-inflammatory innate populations; the resulting TiME retains relatively intact immunostimulatory capacity. AXL^+^ cells were significantly reduced in combination therapy compared to Bem monotherapy (p<0.05), which was not observed in the CSF1R-depleted mice (P<0.05) (**Fig. 3E, Supplementary Fig. S4B**). Bemcentinib has additionally demonstrated macrophage-depleting activity across multiple tumor models in prior published work, consistent with AXL’s role in macrophage survival signaling, providing an additional mechanistic basis for TAM-directed activity beyond gas6:AXL pathway inhibition alone^9,11,12^.There were no significant differences in T cell populations among control and myeloid-depleted animals (**Supplementary Fig. S4C**). CD8^+^ T cell infiltrate did not significantly differ between therapy groups in CSF1R-depleted mice, but increased with Bem therapy in F4/80-depleted animals (**Fig. 3E, Supplementary Fig. S4D**). There were no significant differences in the CD4^+^ T cell infiltrate after either depletion strategy (**Supplementary Fig. S4D-E**).

### AXL expression and function is TAM-context dependent

M1-like and M2-like macrophage differentiation was confirmed by CD86+ and CD206+ expression respectively (**Fig. 5A, Supplementary Fig. S5A-B**). LPS stimulation strongly induced AXL expression in M1-like macrophages, whereas M2-like macrophages were not significantly AXL-positive at baseline (**Fig. 5B**), consistent with AXL’s role in inflammatory macrophage activation. Gas6 reduced AXL expression in M1-like macrophages (p<0.05), while aPD-1 treatment increased AXL expression in M2-like macrophages (**Fig. 5C**). PD-1 expression increased in M1-like macrophages with aPD-1 treatment alone or in combination with Bem (p<0.01 vs. control and Bem groups) (**Fig. 5D**). PD-L1 expression was not significantly altered in either polarization state (**Fig. 5E**). AXL function in efferocytosis was assessed using apoptotic YUMM1.7 cells as macrophage bait; M1-like macrophages demonstrated significantly higher efferocytosis than M2-like macrophages, with no significant modulation by gas6 stimulation or AXL inhibition in either polarization state (**Supplementary Fig. S7A-B, S8A-B**), indicating AXL inhibition does not operate primarily through phagocytic modulation.

**Figure 5.**
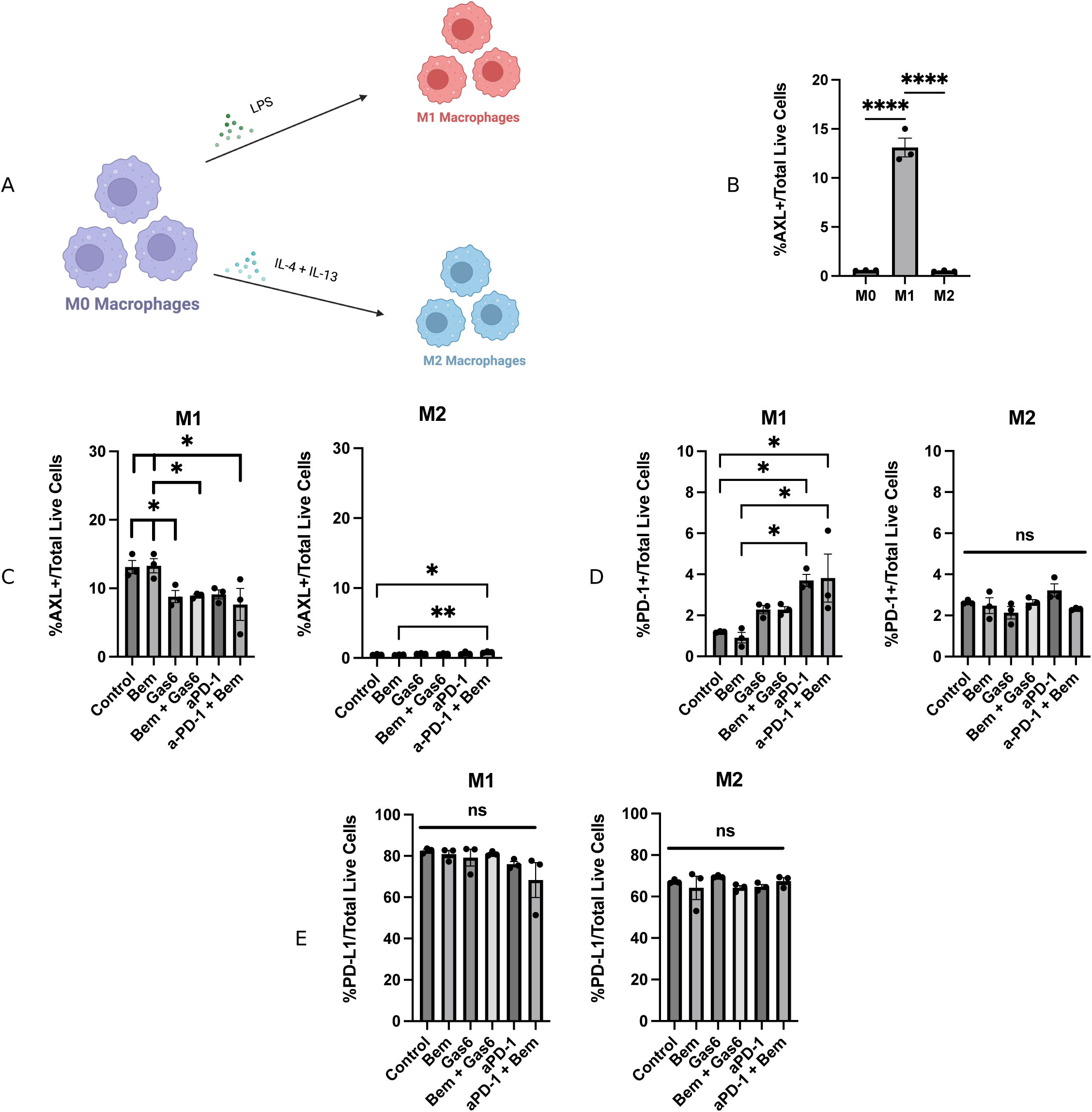
Macrophage polarization impacts AXL expression and aPD-1 synergy. A) Schematic of differentiation protocol. M0 Raw 264.7 macrophages were treated with either LPS or IL-4 + IL-13 for 24 hours to induce M1-like (M1) or M2-like (M2) polarization respectively. B) Baseline AXL expression was measured post-differentiation with significant induction on M1-like polarization. C) Gas6 reduced AXL expression in M1-like but not M2-like macrophages (p<0.05); combination aPD-1 + Bem decreased M1-like but not M2-like macrophages (p<0.05); combination aPD-1+Bem decreased M1-like and increased M2-like AXL expression vs. control or Bem (p<0.05) D) PD-1 expression increased in M1-like macrophages with aPD-1 treatment alone or in combination with bemcentinib (p<0.05 vs control or Bem). No changes were noted in PD-1 expression in M2-like macrophages. E) No changes in baseline PD-L1 expression were noted in M1-like or M2-like macrophages.

### Environmental context impacts AXL and PD-1 synergy

Macrophage paracrine functions were further examined through T cell co-culture and tumor cell migration assays. T cell counts were lower in M2-like vs. M1-like co-culture conditions across treatment groups; combination aPD-1+Bem significantly increased CD8+ T cells vs. gas6 stimulation in M2-like conditions (p<0.05) (**Supplementary Fig. S8A,C-E**). Gas6-stimulated increases in PD-1+CD3+ T cells were reversed by Bem. Secreted factors rather than direct cell contact were the primary driver of macrophage-T cell interactions, with minimal proliferation observed in unconditioned media controls (**Supplementary Fig. S9A-B**). M2-like conditioned media significantly promoted tumor cell migration over M1-like, and Bem reversed this effect (**Supplementary Fig. S9C**).

### AXL-modulation induces changes in the secretome

AXL inhibition with Bem significantly inhibited SK-MEL-24 melanoma cell migration, and M2-like macrophage-conditioned media promoted tumor cell invasion over M1-like conditioned media — an effect reversed by Bem (**Supplementary Fig. S9C**). Macrophage-conditioned media secretome was then analyzed by Luminex assay. Combination therapy reprogrammed the macrophage secretome in a polarization-context-dependent manner. In M2-like macrophages, Bem reversed gas6-driven IL-10 production, confirming AXL involvement in gas6-driven immunosuppressive signaling (**Fig. 6A**). aPD-1 monotherapy drove IL-10 elevation approaching gas6-stimulated levels, while combination therapy reversed this response, synergistically amplifying IL-6 production above all other conditions (p<0.001 vs. aPD-1). Gas6 stimulation elevated CXCL10/IP-10 and CCL5/RANTES in M2-like macrophages, and Bem reversed both (**Fig. 6A**), identifying gas6:AXL signaling as an active chemokine driver in this polarization context. Combination therapy synergistically amplified GM-CSF in M2-like macrophages, and the M2-like secretome heatmap showed a broader immunostimulatory shift with combination therapy across multiple analytes (**Fig. 6C**).

**Figure 6.**
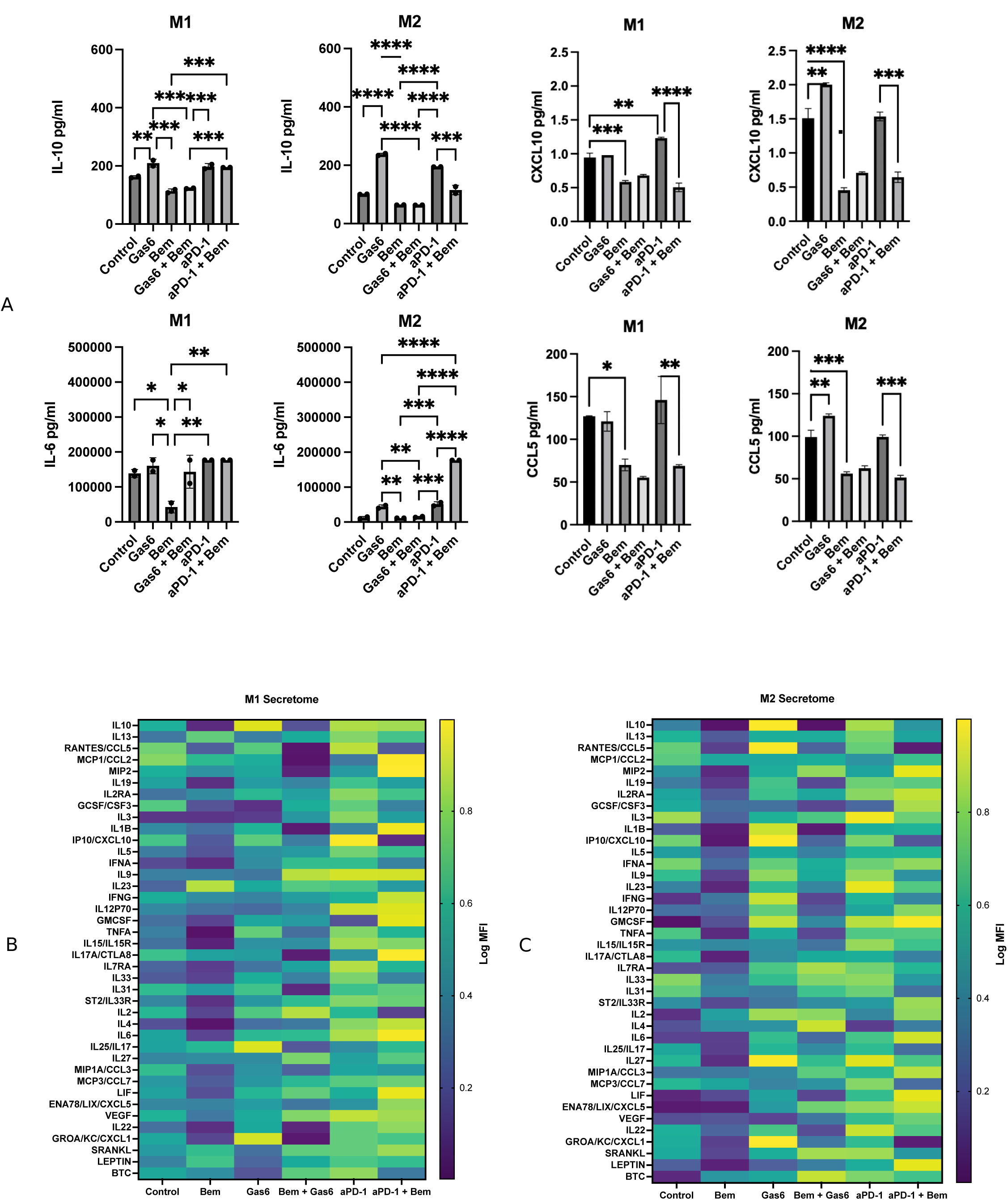
Combination AXL inhibition and anti-PD-1 reprograms macrophage secretome in a polarization-context-dependent manner. Macrophage-conditioned media from M1-like (LPS-stimulated) and M2-like (IL-4/IL-13-stimulated) RAW264.7 macrophages evaluated by Luminex immunoassay under indicated treatment conditions (Control, gas6, Bem, gas6+Bem, aPD-1, aPD-1+Bem). A) Quantitative ELISA of IL-10, IL-6, CXCL10/IP-10, and CCL5/RANTES in M1-like and M2-like macrophages. Values are raw pg/mL from biological triplicates (n=3); y-axis scales are parallelized across M1 and M2 for direct magnitude comparison. Statistical comparisons by one-way ANOVA with Tukey’s post-hoc correction; *p<0.05, **p<0.01, ***p<0.001, ****p<0.0001. B, C) Log min-max normalized heatmap of 40-analyte Luminex panel for M1-like (B) and M2-like (C) macrophages. Heatmap values represent the mean of biological triplicates; normalization applied independently to each panel for visualization of relative cytokine patterns; absolute comparisons require reference to Panel A raw pg/mL data.

In M1-like macrophages, Bem similarly reversed gas6-driven IL-10 production, but combination therapy did not produce the same synergistic IL-6 amplification observed in M2-like conditions (**Fig. 6A**). Bem reduced CCL5 production and attenuated aPD-1-driven elevation of both CXCL10 and CCL5 in combination conditions (**Fig. 6A**). Th1/Th17-associated cytokines were preserved or enhanced in M1-like conditions, confirming selective rather than generalized immunosuppressive effect of AXL inhibition (**Fig. 6B**). Bem alone reduced baseline IL-6, IL-15, CCL5, and CXCL10 in M1-like macrophages, consistent with basal, gas6-independent AXL activity sustaining constitutive cytokine output. IFN-γ and IL-2 did not differ significantly between aPD-1 monotherapy and combination therapy in either polarization state (**Supplementary Fig. S6A**), further supporting a selective rather than broadly immunostimulatory secretome effect. Serum cytokine profiling in tumor-bearing mice showed directionally consistent patterns with in vitro findings, including elevated IL-6 and IL-10 in the combination arm (**Supplementary Fig. S6B-C**). Given n=2 in the combination arm and that serum cytokines reflect systemic rather than intratumoral sources, these observations are presented as supportive systemic concordance rather than primary in vivo mechanistic evidence. Definitive validation through intratumoral cytokine profiling and spatial TAM functional analysis is a specific direction for future work. CD86 and CD206 did not change significantly in either M1-like or M2-like macrophages across treatment conditions (**Fig. 2E**), demonstrating that functional secretome reprogramming occurs without commensurate changes in canonical polarization surface markers, consistent with established evidence of TAM phenotypic plasticity^28^.

## Discussion

Significant interest exists in AXL tyrosine kinase as potential target in multiple tumor types, including immunotherapy-resistant melanoma, but clinical usage of AXL inhibitors has yielded mixed results in early phase trials^39–40^. In particular, the role of AXL as a biomarker for therapeutic selection is continuing to be defined in both tumor cells and the immune microenvironment, with differential expression patterns potentially contributing to the more impressive results with AXL inhibitors in non-small cell lung cancer and triple negative breast cancers^8,39–41^.

While soluble AXL levels in our patient population correlated with advancing melanoma stage, AXL expression in melanoma in TCGA datasets was associated with improved overall survival and aligned with other signatures of inflammation, such as PD-L1 and myeloid infiltrate. From available single cell datasets of treatment-refractory melanoma, we show that AXL is predominantly expressed by TAMs, with only small AXL+ populations of melanoma cells and cancer-associated fibroblasts. This compartment-specific expression pattern may explain the diversity of AXL inhibitor responses across tumor types where AXL+ tumor cell populations predominate, such as pancreatic cancer^42–43^.

AXL inhibition alone significantly depleted TAMs and demonstrated tumor control in ICB-resistant melanoma. We employed CSF1R^44–46^ and F4/80^47–48^ TAM-depletion strategies to selectively target monocyte-derived and tissue-resident TAM compartments. Depletion of the monocyte-derived myeloid compartment abolished AXL inhibitor efficacy, as monotherapy or in combination. Tissue-resident TAM (F4/80) depletion enhanced AXL efficacy as monotherapy and in combination, but not single agent aPD-1 response, indicated that reducing the immunosuppressive TAM barrier is necessary but not sufficient; active reprogramming of the monocyte-derived macrophage secretome by AXL inhibition is required to convert the aPD-1 response from immunosuppressive to pro-inflammatory, making the two interventions mechanistically indispensable as a combination.

These findings converge with clinical data from BGBIL006. In that study, bemcentinib plus pembrolizumab did not improve outcomes in the unselected melanoma efficacy population, and pre-planned tumor cell AXL biomarker analyses did not identify a responding subgroup^18–19^. High AXL expression in inflammatory cells predicted improved pembrolizumab response in exploratory analyses. This is consistent with TAM-predominant AXL expression identified here and supports prospective stratification by TAM composition and functional checkpoint interaction state in future AXL inhibitor trials

Combination therapy reprogrammed the TAM secretome in a polarization-context-dependent manner without altering canonical M1/M2 surface markers. CD86 and CD206 did not change significantly in either polarization state across treatment conditions (**Fig. 2E**), consistent with established evidence that surface phenotyping does not reliably capture TAM functional output^26^. AXL expression showed the same dissociation, with AXL+ TAMs identified across both M1-like and M2-like compartments (**Fig. 1F**), which complicates the use of AXL as a surface classifier of M2-like polarization. These divergent functional profiles support secretome analysis as the more informative mechanistic readout for TAM reprogramming in this system.

AXL inhibition selectively dampened T cell-recruiting chemokine output in M1-like macrophages, both at baseline and under aPD-1 stimulation, without broadly suppressing pro-inflammatory tone. Bem reduced baseline CCL5 and CXCL10 production and attenuated aPD-1-driven elevation of both chemokines in combination conditions (**Fig. 6A**). Th1 and Th17 cytokine production was preserved or enhanced under the same conditions (**Fig. 6B**), confirming a selective rather than generalized immunomodulatory effect. In M2-like macrophages, gas6 stimulation drove significant CCL5 and CXCL10 elevation, both reversed by Bem (**Fig. 6A**), identifying gas6:AXL signaling as an active chemokine driver in this polarization context. Bem dampened baseline IL-6, IL-15, CCL5, and CXCL10 in M1-like macrophages, consistent with ligand-independent basal AXL activity sustaining constitutive cytokine output. In M1-heavy TiMEs, AXL inhibition may limit T cell recruitment by dampening chemokine gradients, while in M2-heavy, AXL-positive, ICB-resistant disease, the combination mechanism predicts meaningful benefit, consistent with the heterogeneous AXL inhibitor results in unselected melanoma populations^41^.

The in vitro M2-like secretome findings provide mechanistic context for the paradoxical checkpoint interaction gain measured by iFRET in vivo. In M2-like macrophages, aPD-1 alone drove immunosuppressive IL-10 elevation approaching gas6-stimulated levels, consistent with evidence that PD-1 engagement on macrophages activates rather than suppresses immunosuppressive signaling in certain tumor contexts. The iFRET efficiency (E*f*) gain in vivo and the M2-like immunosuppressive secretome response to aPD-1 in vitro are therefore convergent observations: aPD-1 engages PD-1 on M2-like TAMs and activates their immunosuppressive program rather than disrupting checkpoint engagement.

Combination therapy resolved both simultaneously: in M2-like macrophages, the combination converted the aPD-1 secretome response from immunosuppressive to pro-inflammatory. In vivo, the combination restored iFRET-measured E*f* to baseline range. PD-L1 expression did not change across treatment groups in either system, confirming a functional rather than expression-level mechanism.

These convergent in vitro and in vivo findings support AXL-driven TAM reprogramming as the mechanism restoring functional checkpoint disruption, and motivate two specific mechanistic questions: how AXL signaling in TAMs stabilizes PD-L1:PD-1 interaction at the molecular level, and which TAM polarization composition predicts ICB resistance and identifies candidates for combination therapy. AXL inhibition did not alter PD-L1 expression in either polarization state, arguing against transcriptional upregulation as the primary driver of checkpoint interaction gain. The mechanism by which AXL signaling in TAMs stabilizes functional PD-L1:PD-1 interaction remains to be defined. Post-translational mechanisms are the more probable candidates: post-translational modification of PD-L1 protein stability, co-receptor complexing at the macrophage surface, or downstream STAT3/PI3K/AKT signaling effects on PD-L1 membrane retention. LRP-1 co-receptor complexing is of particular interest given known co-expression of LRP-1 with AXL on TAMs and emerging evidence for LRP-1 as a modulator of PD-L1 surface availability.

There are several limitations of this study. First, AXL+ non-TAM populations (e.g., NK cells, dendritic cells, and cancer-associated fibroblasts) contribute to immune equilibrium in complex tumor environments and were not independently characterized here. Second, AXL-family receptor redundancy — particularly MerTK and Pros1 — warrants further definition in these models, given evidence here of non-AXL-dependent gas6 functions and ligand-independent basal AXL activity in TAMs. Third, the differential sensitivity of tissue-resident versus monocyte-derived TAMs to AXL targeting requires further definition in patient-derived models. Fourth, the optimal targeting strategy (ligand versus receptor) for both gas6:AXL and PD-L1/PD-1 signaling warrants investigation to maximize ICB response and minimize immune-related toxicities.

Future directions follow directly from the mechanistic gaps identified here. First, **Fun**ctional **o**ncology **map**ping (FuncOmap) is an automated, spatial evolution of the iFRET technique that we have newly integrated with per-pixel multiplex immunofluorescence co-registration^35^ in patient-derived melanoma tissues to determine whether checkpoint interaction gain localizes to TAM-T cell, tumor cell-T cell, or other interfaces. Patient tissue validation is prioritized before further genetic mouse model dissection; while YUMM1.7 AXL knockout models are in development to isolate tumor-intrinsic from macrophage-intrinsic AXL contributions in parallel. Second, a planned cohort of PD-1-refractory melanoma patients will evaluate AXL+ M2-like TAM enrichment as a candidate predictive biomarker, with FuncOmap functional checkpoint interaction state as a co-primary endpoint.

AXL inhibition efficacy is TiME-context dependent. Tailoring AXL blockade to macrophage polarization composition, may maximize therapeutic benefit and advance development of TAM-targeted strategies to reduce immunosuppressive signals in melanoma and other ICB-resistant cancers. A central priority for future work is defining the molecular mechanism by which AXL maintains high PD-L1:PD-1 interaction in ICB-resistant tumors and determining how disruption of this interaction can be optimally achieved in the clinical setting.

## Methods

### Study Design and Models

Single-cell RNA sequencing datasets were obtained from publicly available repositories (GEO, Single Cell Portal, TCGA) and processed using CellRanger (v7.1.0), CellBender (v0.3.0), and Seurat (v4.9.9.9086) as detailed in Supplementary Methods. Soluble AXL was measured by ELISA (R&D Systems, cat# DY154) in blood samples from melanoma patients presenting for surgical excision at UC Davis Comprehensive Cancer Center (2016–2020).

### In vivo

Four- to six-week-old female C57/Bl6 mice were inoculated subcutaneously with 1×10L YUMM1.7 melanoma cells and randomized to receive saline control, warfarin (500 nM in drinking water), bemcentinib (50 mg/kg daily by oral gavage, cat# HY-15150, MCE), anti-PD-1 (1 mg/kg 3x/week intraperitoneally, Invivomab cat# BE0146R00, Bio X Cell), or combination for 14–21 days until control tumors reached 1,500 mm³. For myeloid depletion studies, anti-CSF1R (clone AFS98, 2 mg per mouse) or anti-F4/80 (clone CI:A3-1, 0.5 mg per mouse) antibodies were administered by intraperitoneal injection every 3 days beginning 7 days prior to tumor inoculation and continuing throughout the treatment period. All animal studies were conducted under APLAC-approved protocols at Stanford Cancer Institute. Full in vivo protocols are provided in Supplementary Methods.

### In vitro

M1-like and M2-like macrophages were derived from RAW264.7 cells (ATCC) by stimulation with LPS (100 ng/mL) or IL-4/IL-13 (20 ng/mL each) respectively for 24 hours. Polarization was confirmed by flow cytometry (CD86 for M1-like, CD206 for M2-like) prior to treatment initiation. Macrophage-conditioned media was collected after 24 hours of treatment and analyzed by Luminex immunoassay; serum cytokine profiling used a 48-plex Procarta panel (cat# EPX480-20834-901) with n=2 samples available in the combination arm due to variable serum recovery. Treatments included gas6 (100 ng/mL, cat# 67202S, Cell Signaling), bemcentinib (0.5 µM), and anti-PD-1 (0.5 nmol/L). Full in vitro protocols are provided in Supplementary Methods. Functional PD-L1:PD-1 checkpoint interactions were quantified by immune Förster Resonance Energy Transfer (iFRET) using the FuncOmap automated spatial platform as previously described32-34. Banafshë Larijani is co-inventor of patents held by the Francis Crick Institute (US 10,578,620 B2; PCT/EP2018/062719) relating to methods for detecting molecules and cell-cell interactions used in this platform.

### Statistical Analysis

The data were examined using GraphPad Prism 10 for Windows (GraphPad Software; www.graphpad.com). Findings are presented as mean ± standard error of the mean (SEM) or SD as indicated. Two-group comparisons used unpaired t-tests. Multi-group comparisons used one-way ANOVA with Tukey’s post-hoc correction. Individual Luminex analytes were analyzed by one-way ANOVA with Tukey’s post-hoc correction; no cross-analyte multiple comparison correction was applied, and the full panel is designated as hypothesis-generating. In vivo CSF1R depletion timepoint analyses were conducted as independent per-timepoint ANOVAs to account for accelerated tumor growth in that arm. p<0.05 was considered statistically significant.

## Acknowledgements

We thank the generous support of the John and Marva Warnock Scholar Fund, UC Davis Immuno-Oncology Innovation Award, and the Stanford Department of Surgery in the completion of this work. We thank M.Usman Ahmad for significant administrative support and Hsiu “Sherry” Hsu for thoughtful input.

