## Supplemental Methods for "AXL inhibition reprograms tumor-associated macrophages to restore immune checkpoint blockade efficacy in a polarization-context-dependent manner"

**Supplementary Methods**

**Single cell sequencing**

Data Extraction: We collected five single-cell RNA sequencing (scRNA-seq) datasets and two bulk RNA sequencing datasets for humans from publicly available repositories, including Gene Expression Omnibus (GEO), Single Cell Portal, and The Cancer Genome Atlas (TCGA). For the 10X Genomics data, Raw sequencing files were obtained using SRA-Toolkit (v3.0.2) via the fastq-dump utility. After obtaining the Raw sequencing data, gene expression (GEX) matrices were generated through a comprehensive pipeline. First, the data were demultiplexed to separate individual cell barcodes from sequencing reads. Next, barcode processing was performed to filter out low-quality or ambiguous barcodes, ensuring accurate assignment of reads to specific cells. The processed reads were then aligned to the human reference genome Hg38 using a highly optimized alignment algorithm. Gene quantification was carried out, whereby aligned reads were counted to generate a matrix of gene expression levels across all cells. All steps in this pipeline were performed using the 10X Genomics CellRanger software (v7.1.0). For datasets generated using the SMART-Seq2 protocol, GEX matrices were directly downloaded from the Gene Expression Omnibus (GEO) and Single Cell Portal repositories.

**Single-cell RNA data filtering and normalization**

To enhance the accuracy of gene expression estimates, we utilized the `remove-background` function in CellBender (version 0.3.0) for each sample. CellBender utilizes a deep learning-based approach to model and remove technical artifacts, such as background noise and ambient RNA, which can contaminate single-cell RNA sequencing data. By estimating the true biological signal from the Raw gene-by-cell count data, CellBender generates ambient-corrected count matrices that provide a more accurate reflection of the cellular transcriptome. These corrected matrices were then imported into R (version 4.2.2) and converted into Seurat objects using Seurat (version 4.9.9.9086). We applied stringent cell selection criteria to ensure data quality: cells were retained if they exhibited less than 10% or up to 20% mitochondrial reads, expressed between 3,500 and 7,500 genes and had unique molecular identifier (UMI) counts ranging from 500 to 60,000. Next, doublet cells were identified using Scrublet (version 0.2.3), which computes doublet scores based on the expected doublet rate, preset at 10%. Cells flagged as doublets by Scrublet with the default parameters were excluded from further analysis.

For datasets generated using the SMART-Seq2 protocol, we incorporated pre-filtered datasets sourced from public repositories, including GEO and Single Cell Portal. After data filtering, the gene-by-cell expression matrices were merged using the `merge` function. We then proceeded with pre-processing using SCTransform (version 0.4.0), which incorporates a variance-stabilizing transformation (VST) method. SCTransform replaces conventional normalization and scaling procedures with a regularized negative binomial regression model, effectively controlling for technical noise. This method adjusts gene expression measurements by modeling unwanted variation and emphasizing biological variability.

**Enzyme-linked immunosorbent assay (ELISA)**

Our group measured soluble AXL by solid phase sandwich ELISAs according to the manufacturer’s protocol ( cat# DY154 R&D Systems, Minneapolis, USA) in blood samples of all melanoma patients at the UC Davis Comprehensive Cancer Center presenting for surgical excision of melanoma from 2016-2020.

**In vivo experiments**

**Animal studies**

All animals were housed in a pathogen-free facility with 24-hour access to food and water. Experiments were approved by, and conducted in accordance with, an APLAC approved protocol at Stanford Cancer Institute for therapeutic studies with warfarin and bemcentinib. Bemcentinib studies were repeated for a total of three independent experiments (5 animals per group, total of 75 animals) to serve as controls for the myeloid depletion models. Four- to six-week-old female C57/Bl6 mice were obtained from Jackson Laboratories. A total of 1×10^6^ Yumm1.7, Yummer1.7, and B16F10 melanoma cells, purchased from the ATCC, were injected orthotopically as described^57^. Mice were randomized to receive 0.5 mg/L warfarin (500 nM) once tumors were measured at caliper volume of 100mm^3^. In subsequent experiments, mice were treated with saline controls, bemcentinb (50 mg/kg) (cat# HY-15150, MCE), Invivomab aPD-1 (1 mg/kg) (cat# BE0146R00, Bio X Cell) or combination as described. Bemcentinib was administered daily by oral gavage and aPD-1 by intraperitoneal injection 3x/week. Therapy was administered for 14-21 days until control tumors reached 1,500 mm^3^.

For myeloid depletion studies, anti-CSF1R (clone AFS98, 2 mg per mouse) or anti-F4/80 (clone CI:A3-1, 0.5 mg per mouse) antibodies were administered by intraperitoneal injection every 3 days beginning 7 days (anti-CSF1R) or 3 days (anti-F4/80) prior to tumor inoculation and continuing throughout the treatment period.

Terminal serum was collected by cardiac puncture at time of sacrifice. Serum recovery was variable across animals, particularly in later-timepoint combination arm animals. The combination arm (aPD-1 + Bem) yielded n=2 serum samples with sufficient volume for Luminex analysis; all other groups yielded n=3. Combination arm observations in Supplementary Fig. S6 are exploratory and directional; formal statistical comparisons are reported only for groups with n≥3.

**Luminex assay**

Mouse serum samples were collected on day 18, at time of sacrifice. Mouse 48-plex Procarta kits (EPX480-20834-901) were purchased from Thermo-Fisher/Life Technologies and used according to the manufacturer’s recommendations with modifications as described. This assay was also used to analyze the secreted cytokines and chemokines present in our in vitro T cell, macrophage, and tumor cell coculture. In brief, beads were added to a 96-well plate and washed in a BioTek ELx405 washer. Samples were added to the plate containing the mixed antibody-linked beads and incubated overnight at 4°C with shaking. Cold (4℃) and room temperature incubation steps were performed on an orbital shaker at 500-600 rpm. Following the overnight incubation, plates were washed in a BioTek ELx405 washer and biotinylated detection antibody was added for 60 minutes at room temperature with shaking. The plate was washed, and streptavidin-PE was added for 30 minutes at room temperature, after which the plate was washed and reading buffer added to the wells. Each sample was measured in duplicate. Plates were read using a Luminex 200 or a FM3D FlexMap instrument with a lower bound of 50 beads per sample per cytokine. Custom Assay Chex control beads (Radix BioSolutions) were added to all wells. The data was then processed in Microsoft Excel using logarithmic min-max normalization.

**Luminex data analysis.** Quantitative cytokine and chemokine levels are reported as raw pg/mL for individual analytes. These values are used for all statistical comparisons. For heatmap visualization, log min-max normalization was applied independently to each panel to enable visualization of relative cytokine patterns across conditions within each experimental context. Normalization is applied for visualization only; quantitative comparisons use unnormalized pg/mL values. In vitro heatmap values represent the mean of biological triplicates (n=3); the in vivo heatmap displays individual animal replicates to convey biological variance.

**Flow cytometry**

Single-cell suspensions were prepared from sacrificed mice tumor samples and stained with fluorescently-labeled antibodies according to the manufacturer's instructions. Cell sorting/flow cytometry analysis for this project was performed on instruments in the Stanford Shared FACS Facility (RRID:  SCR_017788) with appropriate gating strategy to identify specific cell populations. Data analysis was performed using FlowJo software (Tree Star) to quantify and analyze cell populations. The list of antibodies and kits used includes the following: Zombie Violet™ Fixable Viability Kit (cat# 423113), PE/Cyanine7 anti-mouse CD3 (cat# 100220), PE anti-mouse CD4 (cat# 100408), FITC anti-mouse CD8a (cat# 100706), PE anti-mouse CD68 Antibody (cat# 137014), Brilliant Violet 650™ anti-mouse CD86 (cat# 105035), Brilliant Violet 785™ anti-mouse CD206 (MMR) (cat# 141729), PE/Cyanine7 anti-mouse CD163 Antibody (cat# 155320), FITC anti-mouse F4/80 Antibody (cat# 123108), Brilliant Violet 570™ anti-mouse/human CD11b (cat# 101233), Brilliant Violet 750™ anti-mouse CD11c (cat# 117357), Alexa Fluor® 700 anti-mouse CD68 (cat# 137025), PE anti-mouse CD279 (PD-1) (cat# 114117), and PE/Cyanine7 anti-mouse CD274 (B7-H1, PD-L1) (cat# 124314) purchased from BioLegend (BioLegend Way, San Diego, CA 92121 United States) and antigen-presenting cell (APC)/AXL Monoclonal Antibody (MAXL8DS) (cat# 17-1084-82) from eBioscience™, Thermofisher.

**Two-site Antibody labelling**

The tissue microarrays (TMAs) were constructed in collaboration with BioIVT. Tissue sections were reviewed under dermatopathological guidance for the selection of appropriate regions for sampling. Core samples (2 mm) were obtained from formalin-fixed paraffin-embedded (FFPE) blocks of the tumor core, tumor:normal skin interface, post-therapy tumor bed, and DSLN. TMAs underwent an antigen retrieval process using the Envision Flex retrieval solution, pH 9. The Dako PT-Link system was utilized, for which the slides were heated to 95^°^C for 20 minutes. Using a PAP pen, an aqueous-repelling border was outlined around each tissue fragment. Pierce endogenous peroxidase suppressor was then applied to each specimen, and the slides were left to incubate for 30 minutes at 21^°^C room in a humid-controlled environment. The samples underwent two washes with phosphate-buffered saline (PBS) before being incubated for an hour at room temperature with 3% BSA (10 mg/ml). The donor-only slides were incubated with aPD-1 (at a dilution of 1:100). The donor-acceptor slides were treated with the following primary antibodies: aPD-1 (1:100) and anti-PD-L1 (1:500). Primary antibodies were incubated overnight at 4°C. Samples were washed with 0.02% PBS-tween (PBST). The samples were then treated with secondary F(ab’)2 fragments. F(ab’)2- Atto 488 (at a dilution of 1:100) was introduced to the donor-only slides. The donor-acceptor slides received both F(ab’)2-Atto 488 (1:100) and F(ab’)2- horseradish peroxidase (HRP) (1:200). These samples were incubated for 2 hours in the dark at room temperature in a humidified container. After the incubation period, slides were washed with 0.02% PBST. The donor-only slides were mounted with 1 drop of Prolong Glass antifade mount. The donor-acceptor slides were subjected to TSA. The purpose of TSA is to amplify the acceptor labeling, which increases the signal-to-noise ratio and enhances the resonance energy transfer. This procedure is described in detail in Magraner Sanchez *et al.* (2020).^58^ The antibodies were labeled with species-specific F(ab’)2 fragments which were conjugated to Atto 488 (donor chromophore, used to label the receptor primary antibody) or HRP (used to label the ligand primary antibody). TSA was used to conjugate the acceptor chromophore (Alexa 594) to the HRP labeling the ligand.

Banafshé Larijani is the co-inventor of the following patents (Held by the Francis Crick Institute, UK): Patent - US 10,578,620 B2- Methods for detecting molecules in a sample. Patent - PCT/EP2018/062719-Kits, methods, and their uses for detecting cell-cell interactions.

**Time-Resolved Immune Förster-Resonance-Energy Transfer (iFRET) Determined by Frequency-Domain FLIM.**

The quantitative molecular imaging platform utilizes a custom-made semi-automated frequency-domain FLIM (Lambert Instruments). The first slide (donor only) is excited by a modulated (40MHz) diode 473nm laser, and the lifetime of the donor alone recorded. The second slide waw excited by the diode modulated 473nm laser and lifetime of the donor in the presence of the acceptor recorded. The reduction of donor lifetime (caused by resonance energy transfer) due to the presence of the acceptor reports on distances of 1-10nm and therefore acts as a “spectroscopic ruler” enabling to quantify receptor-ligand protein interactions.

We identify the coincidence regions where both the donor and acceptor are observed and their Ef values were calculated and mapped on to the expression levels of PD1.

**Photophysical parameters for quantification of protein interactions.**

We calculate lifetime image of the donor in the presence of an acceptor (𝛕_DA_) and a lifetime image of the donor (𝛕_D_), followed by calculation of reduction of 𝛕_DA_ compared to 𝛕_D_, which is reflected in a parameter called FRET-efficiency (*Ef*):

$Ef$  = [1 − <t _DA_>/<t_D_>]x100	**Eq-1**

FRET-efficiency was calculated as an average for each coincident region. 𝐸𝑓 is directly related to the distance between the donor and acceptor fluorophores (Atto 488 and Alexa 594) (Eq-2), where r is the distance between Atto 488 and Alexa 594 in these experiments. 'R_0_' the Förster radius in this case is 5.83 nm and it is the distance whereat transfer efficiency is 50%.

$Ef=\frac{R_{0}^{6}}{R_{0}^{6}+r^{6}}$ **Eq-2**

$r= \sqrt[6]{R_{0}^{6}*\frac{(1-Ef)}{Ef}}$   **Eq-3**

**Statistical analysis of FRET**

Violin plots were used to visualize the distribution of Ef values across tissue regions and samples. The violin plots are generated using all the coincident images. The median of the global E*f* distribution was used as a descriptive metric to represent regions with the highest PD1/PD-L1 interaction states across tissues. Following this, the Mann-Whitney U test was utilized to statistically analyze and compare the FRET efficiencies between groups. A p-value of <0.05 was considered statistically significant.

***In vitro* experiments**

**Western blotting**

Cells were washed with PBS on ice and collected by scraping in cold PBS. The cell pellet was lysed in RIPA buffer (cat# 786-489, G-Biosciences, USA) in presence of Halt™ protease and phosphatase inhibitors (cat#78440, ThermoFisher, USA). The protein concentration was determined using the Bio-rad Protein Assay (Bio-Rad, USA) and an equal amount of proteins (40 µg) was loaded on pre-casted SDS Midi protein gel (cat# 5678123, Bio-Rad, USA). The protein was transferred from the gel to PVDF membrane (cat# 1704157, Bio-Rad, USA) using Transblot semi-dry (Bio-Rad, USA). PVDF membrane was blocked in 5% blotting grade dry milk in PBS with 0.1% Tween (PBST) for 1 h at room temperature and then membrane was washed 2 times with PBST (0.05% tween 20 in 1X PBS) for 10min each. Then membrane was incubated with Total AXL primary antibody diluted (1:100, cat# MAB8541, Monoclonal, R&D, USA) in 5% BSA in PBST overnight at 4 °C. The next day membrane was washed 2 times with PBST (0.05% tween 20 in 1X PBS) for 10min each and incubated with the appropriate secondary antibody (1:1000, cat# A18745, HRP- conjugated anti-rat, Invitrogen, USA) for 1 h at room temperature. The bands were then visualized using enhanced ECL reagent (cat#170-506, Bio-Rad, USA). The analysis was carried out using Chemi Doc (BioRad, Image Lab 5.1).

**Macrophage differentiation**

M1 and M2 macrophages were derived from Raw 264.7 mouse macrophages (ATCC). M1 macrophages were activated with 100 ng/mL LPS (cat# 2939, Tocris) for 24 hours, while M2 macrophages were polarized with 20 ng/mL of IL-4 (cat# 200-04, Peprotech) and IL-13 (cat# 200-13, Peprotech) for 24 hours. In addition, Flow cytometry analysis was done to measure specific macrophage markers including CD86, CD206, as well as checkpoint molecules PD-1 and PD-L1 utilizing our previously described panel of immunofluorescent antibodies.

**In Vitro Migration Assay**

1x10^6^ Yumm1.7 or SK-MEL-24 were cultured in RPMI 1640 media (cat# 11875093, Thermofisher) with 10% FBS (cat# MT35011CV, Corning) in each well of 6-well cell culture plates until they reached at least 70% confluency. After confluency was reached, media from each well was aspirated and gas6 (cat# 67202S, Cell Signaling) or warfarin (bemcentinib was utilized in the SK-MEL-24 migration assay) were added in 1% FBS RPMI 1640 media. In addition for the SK-MEL-24 experiment, the culture media was substituted Raw cell-conditioned media with or without addition of gas6 or bemcentinib. 100um scratch was performed and migration measured at time 0 and 24 hours^57^. Images taken using a Zeiss Axio Observer fluorescent microscope. Images compiled and analyzed with ImageJ using the measuring tool to determine the width of the wound at 5 points along the wound and plotted using Graphpad software as % closure. All assays were performed in triplicate.

**Efferocytosis Assay**

M1-like and M2-like macrophages were derived from Raw 264.7 mouse macrophages (ATCC) through differentiation protocols utilized in our in vitro assays. The macrophages were then labeled with Cytotell blue dye (cat #601770, Efferocytosis Kit, Cayman Chemicals). Yumm 1.7 melanoma cells (ATCC) were cultured in RPMI 1640 supplemented with 10% FBS. Apoptotic Yumm 1.7 cells were induced by treating cells with 5 μM staurosporine (cat# S1421, SelleckChem) for 6 hours, after which cells were labeled with CFSE green-fluorescent dye (cat #601770, Efferocytosis Kit, Cayman chemicals). Co-culture experiments were performed in duplicate with each experiment initiated by seeding 1x10^6^ macrophages per well in 5 wells of 6-well plates: one M0, two M1, and two M2 wells. This was followed by the addition of labeled apoptotic cells at a ratio of 5:1 (apoptotic cells:macrophages). At this step, bemcentinib (0.5 μM) and gas6 (100 ng/ml) were added to cultures. After a 2-hour incubation at 37°C, the co-culture samples were fixed in 4% PFA for immunofluorescence staining using CD86 (cat# 105035, Biolegend) and CD206 (cat# 141729, Biolegend). Flow cytometry analysis was employed to quantify efferocytosis, measuring the percentage of macrophages that had engulfed apoptotic cells through the colocalization of Cytotel blue and CFSE dyes.

**T Cell Interaction Assay**

Raw 264.7 mouse macrophages (ATCC) were differentiated into either M1 or M2 subtypes using our established in vitro protocol for 24 hours. On the next day, we initiated the apoptosis of Yumm 1.7 mouse melanoma cells (ATCC) using 5 μM staurosporine (cat# S1421, SelleckChem) for 8 hours. At the conclusion of apoptosis, the Yumm 1.7 melanoma cells were added to the differentiated macrophages at a concentration of 4 x 105 cells per well and were incubated overnight. In this step, we also added drugs of interest which included 0.5 uM Bemcentinib (cat# HY-15150, MCE), 100 ng/ml gas6 (cat# 67202S, Cell Signaling), and 0.5 nmoles/L aPD-1 (cat# BE0146R00, Bio X Cell), in triplicate plating conditions. In parallel, we collected ten spleens from healthy euthanized mice. The spleens were processed into single cells using mechanical digestion, and T cells were isolated using the Dynabeads Untouched Mouse T Cell kit (cat# 11413D, ThermoFisher Scientific). After isolation, the T cells were labeled with cellTrace carboxyfluorescein succinimidyl ester (CFSE) (cat# C34554, Invitrogen) and were incubated at 37℃ in a flask with RPMI 1640 media (cat# 11875093, ThermoFisher Scientific), 10% FBS (cat# MT35011CV, Corning), 1% penicillin/streptomycin (cat# 15140122, Gibco), 50 μM β-mercaptoethanol (cat# 21985023, Thermofisher) and 2 mM L-glutamine (cat# 25030081, Gibco) to expand them overnight. After a 24-hour incubation, T cells were introduced to the coculture system involving tumor bait-exposed macrophages by adding 1 million T cells per well. After an additional 24-hour incubation, the final step of the experiment involved the analysis of T cells by flow cytometry analyzing CD3 (cat# 100220, BioLegend), CD4 (cat# 100408, BioLegend), CD8 (cat# 100706, BioLegend), and PD-1 (cat# 114117, BioLegend). In addition, the media from the triculture system was collected, and cytokine profiles were analyzed via Luminex as detailed previously.
