## Supplementary Figures for "AXL inhibition reprograms tumor-associated macrophages to restore immune checkpoint blockade efficacy in a polarization-context-dependent manner"

Supplementary Figure 1

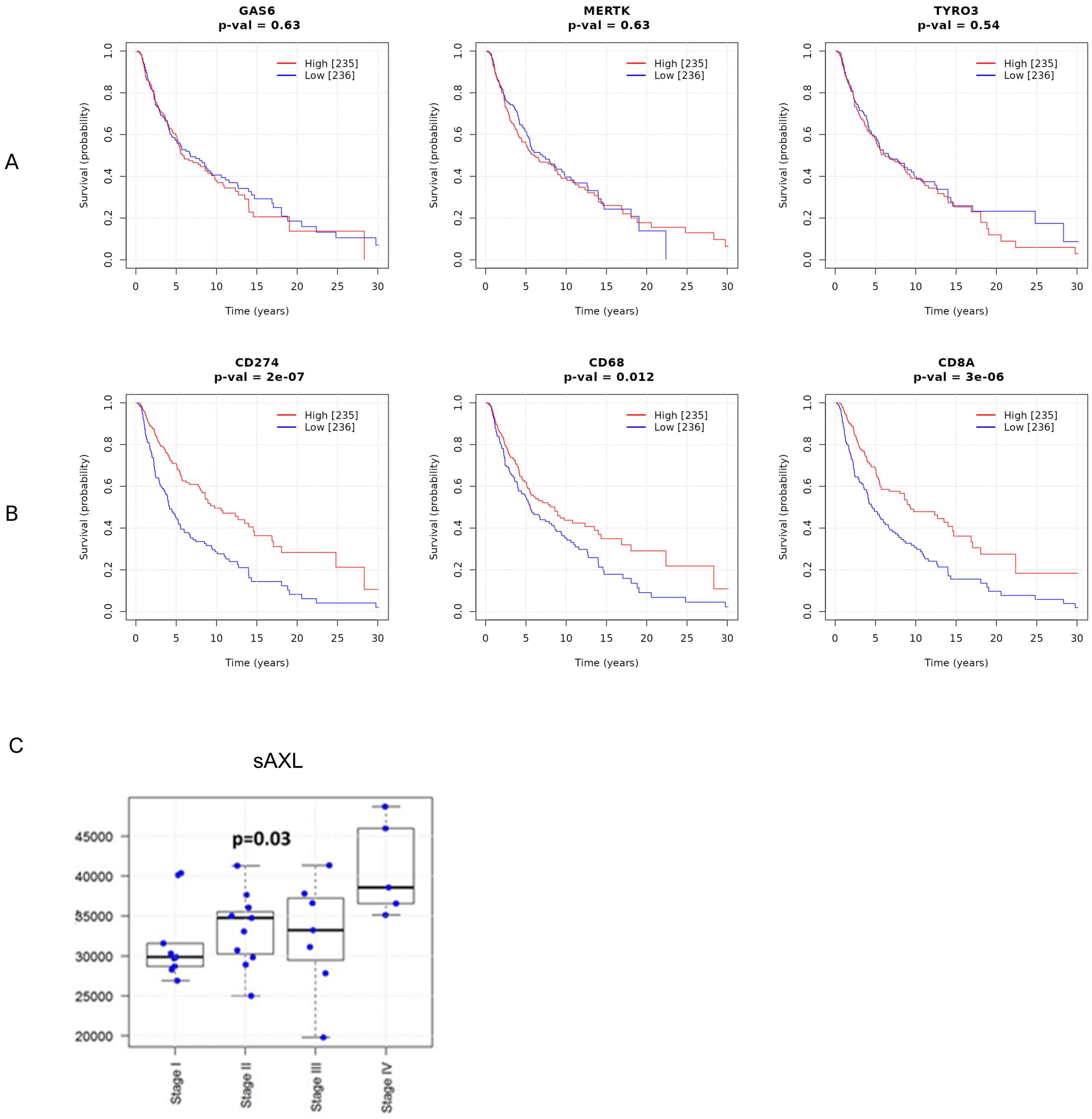

**Supplementary Figure S1. Clinical correlates of AXL tyrosine kinase family receptors and ligands with survival.** A) Kaplan-Meier survival curves created from TCGA-SKCM data demonstrate no correlation with AXL ligand gas6 or other AXL family kinases (Tyro3, MerTK). B) An overall survival benefit is seen in correlation with inflammatory markers and cytotoxic T cells (CD274/PD-L1, CD68, CD8A shown here). C) Soluble AXL (sAXL) is elevated in patients with Stage IV melanoma relative to earlier stage disease, consistent with active pathway engagement in advanced disease.

Supplementary Figure 2

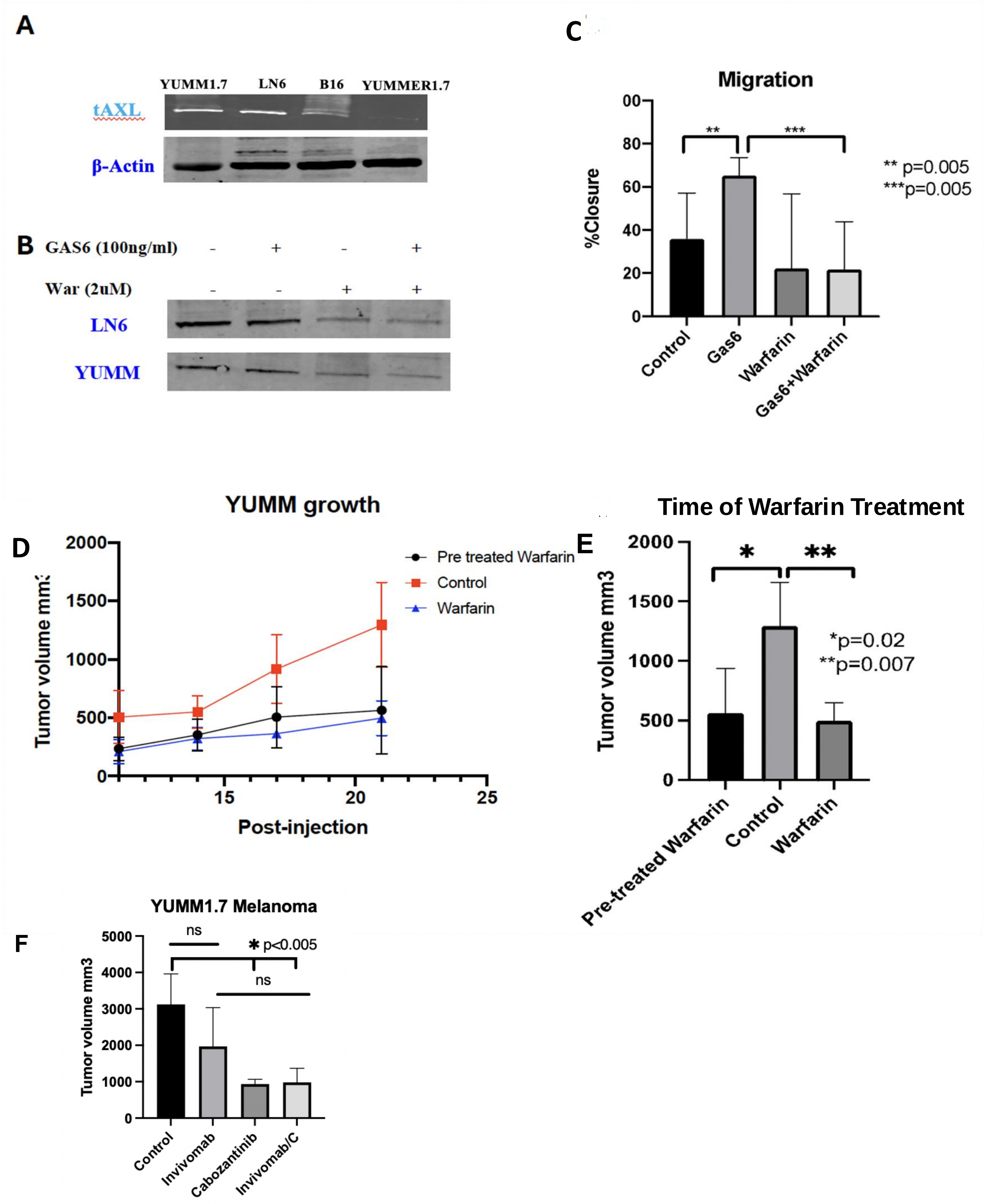

**Supplementary Figure S2**. AXL targeting in anti-PD-1-resistant melanoma. A) Melanoma cell lines were screened for baseline AXL expression in vitro, revealing higher expression in anti-PD-1-resistant models over AXL-low, PD-1 sensitive Yummer1.7 with B) inhibition of p-AXL by warfarin exposure in vitro. C) Migration of Yumm1.7 cells was significantly stimulated by gas6 ($p=0.02$) and inhibited by warfarin 2 µM ($p=0.007$). D-E) Mice (n=6/group) were pretreated with warfarin for 7 days prior to subcutaneous injection of Yumm1.7 tumor cells versus randomized to therapy with warfarin after establishment of tumors (>150 mm3). No differences in tumor size were observed between warfarin administration before or after tumor establishment with both treatment groups demonstrating significant tumor growth reduction ($p<0.05$). F) Similar tumor control was observed with cabozantinib, a multi-kinase inhibitor with AXL inhibitory activity (cMET, VEGFR2, RET, AXL among other kinases), consistent with AXL pathway engagement as the relevant mechanism. Results are supportive of AXL pathway involvement but cannot be attributed exclusively to AXL inhibition. Anti-PD-1 antibody (Invivomab, Bio X Cell cat# BE0146R00) is referred to as aPD-1 throughout.

Supplementary Figure 3

**A**

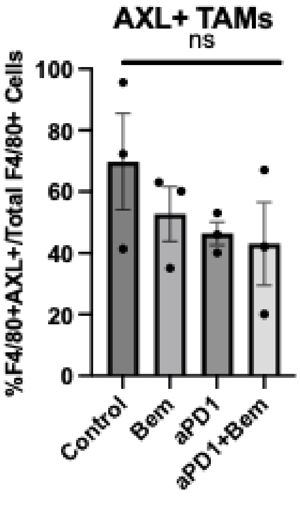


**B**

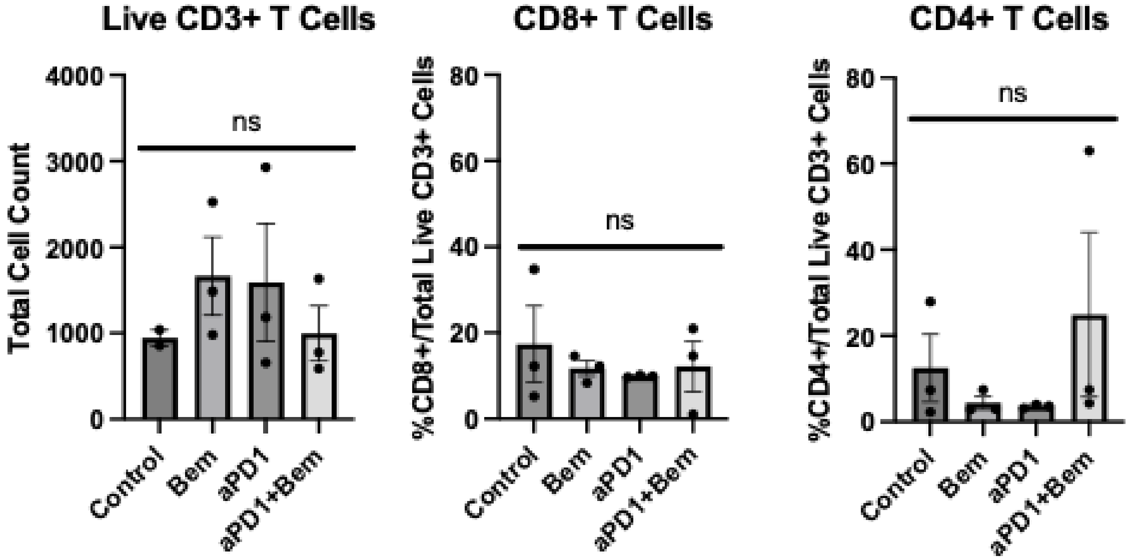

**Supplementary Figure S3. Macrophage and T cell composition of tumors.** A) AXL+ TAM distribution was not significantly altered between treatment groups in YUMM1.7 tumors. B) No significant differences were observed among treatments in total CD3+ T cells, CD8+CD3+ T cells, or CD4+CD3+ T cells.

Supplementary Figure 4

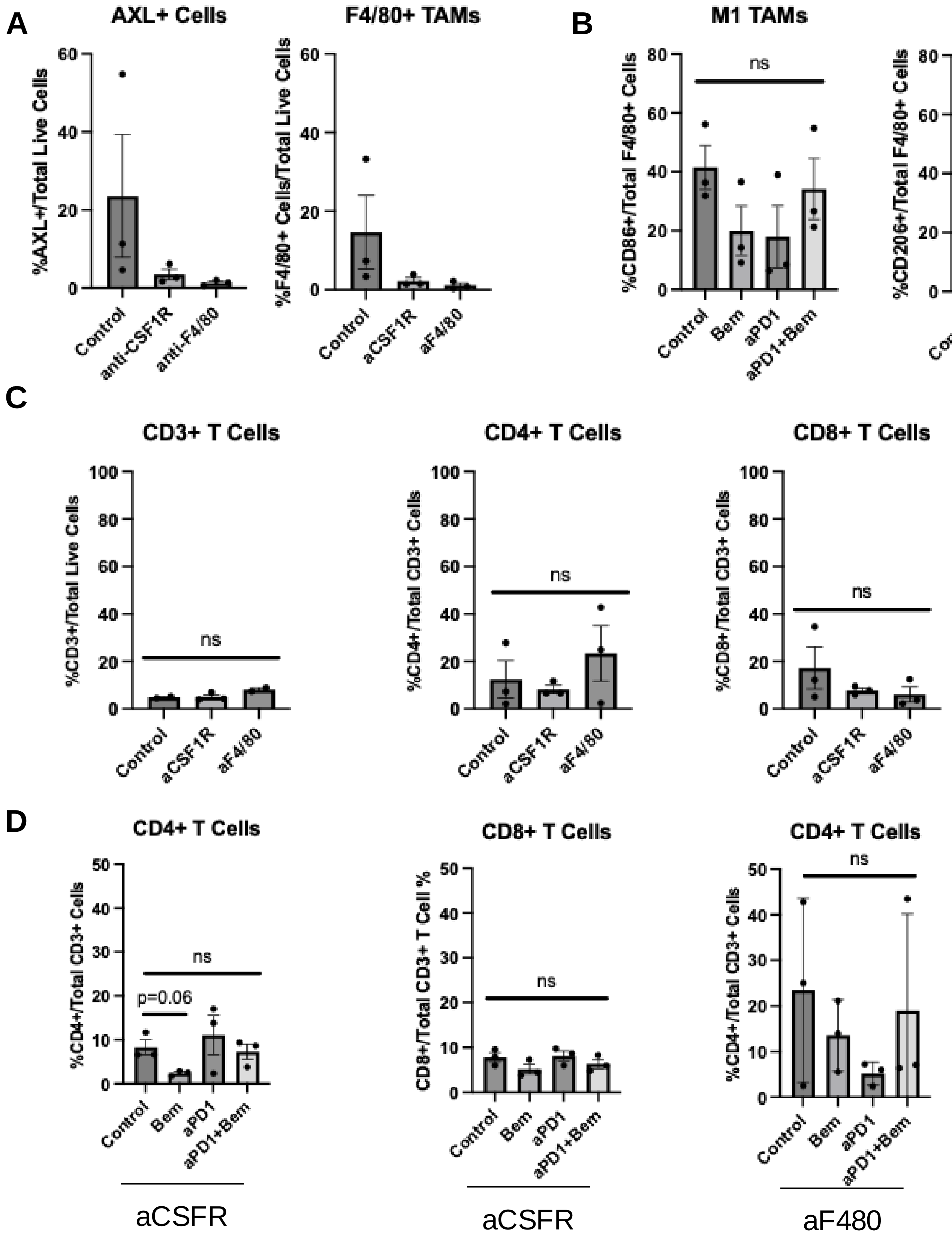

**Supplementary Figure S4. TiME composition of tumors with CSF1R or F4/80 depletion treated with anti-AXL and aPD-1.** A) Anti-CSF1R and anti-F4/80 treated mice were depleted of CSF1R+ monocyte-derived myeloid cells and F4/80+ tissue-resident macrophages, respectively, confirming depletion efficiency. B) No significant changes in M1-like or M2-like TAM compositions in anti-CSF1R depleted mice. C) No significant differences were observed among mice with or without myeloid depletion in total CD3+, CD8+CD3+, or CD4+CD3+ T cells. D) CD8+ T cell infiltrate was not altered by aPD-1 therapy or bemcentinib in CSF1R depleted mice. Bemcentinib reduced CD4+ T cells in CSF1R depleted mice but was below statistical significance ($p<0.06$). No differences were significant in CD4+ T cells with anti-F4/80 depletion.

Supplementary Figure 5

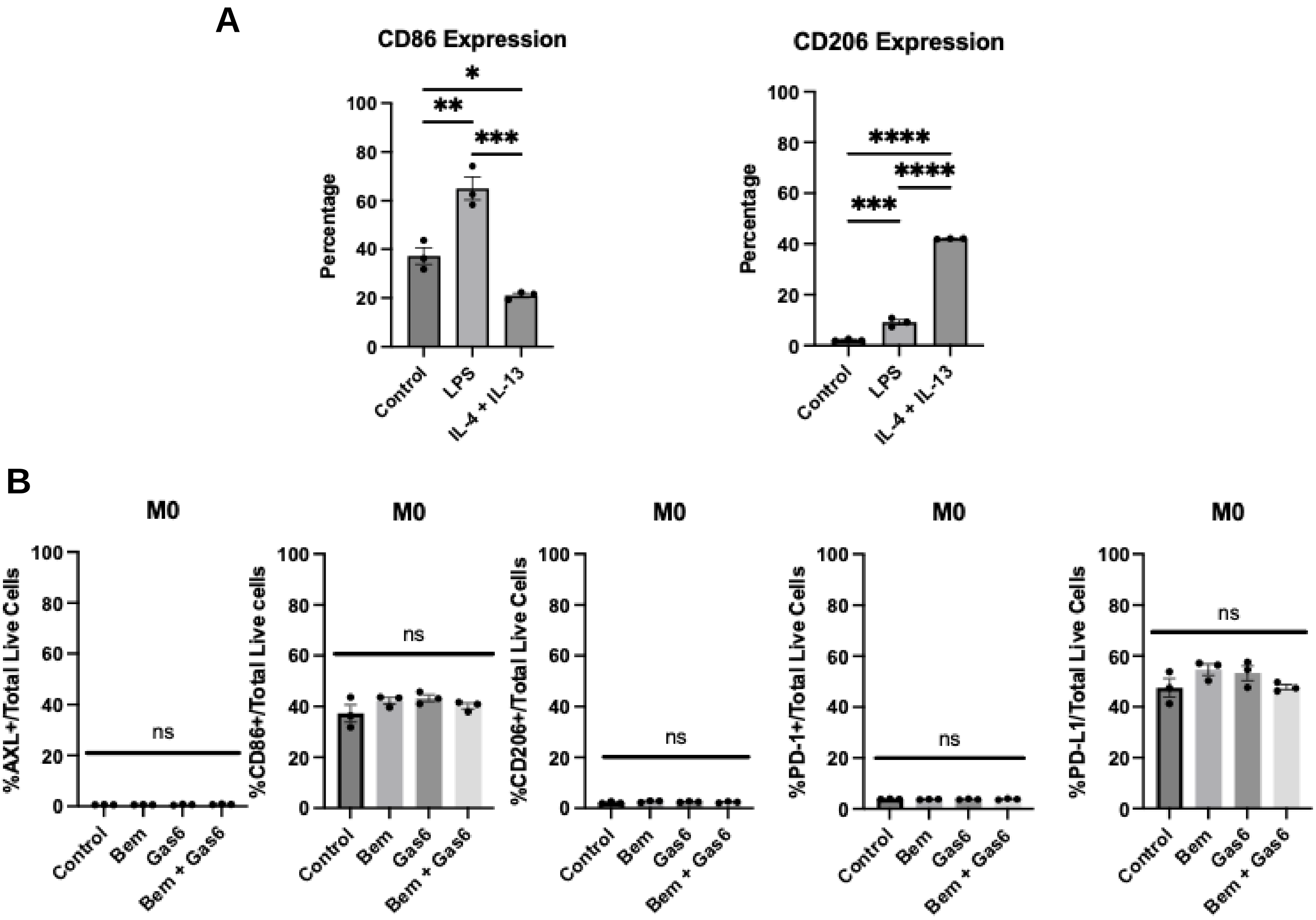

**Supplementary Figure S5**. Impact AXL modulation on Raw macrophage polarity. A) LPS strongly induces M1-like polarization (CD86+), IL-4 and IL-13 induce M2-like polarization (CD206). B) AXL, CD86, CD206, PD-1, and PD-L1 expression is not affected by bemcentinib or gas6 alone in nonpolarized macrophages.

Supplementary Figure 6

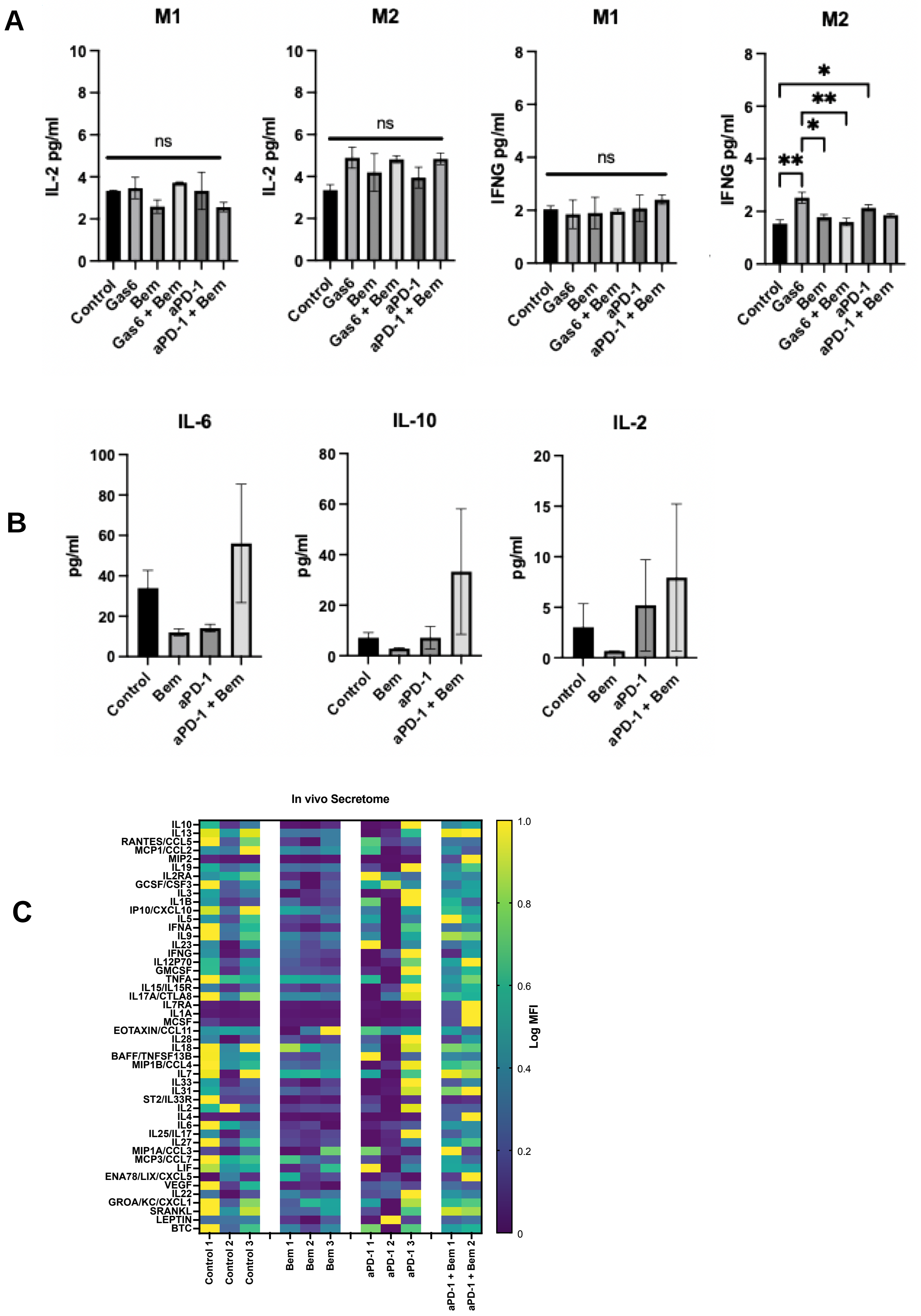

**Supplementary Figure S6. In vitro cytokine secretome and in vivo serum cytokine profiling.** A) IFNG and IL-2 production in M1-like (LPS-stimulated) and M2-like (IL-4/IL-13-stimulated) RAW264.7 macrophages under indicated treatment conditions by Luminex ELISA; values are raw pg/mL from biological triplicates (n=3). IFNG and IL-2 did not differ significantly between aPD-1 monotherapy and combination therapy in either polarization state. B) Quantitative serum ELISA of IL-6, IL-10, and IL-2 in YUMM1.7 tumor-bearing mice under indicated therapy conditions at time of sacrifice. Sample availability in the combination arm (aPD-1 + bemcentinib) was limited to n=2 animals due to insufficient serum recovery from moribund animals at late timepoints; this precludes statistical significance claims for the combination condition. Directional patterns are consistent with in vitro M1-like and M2-like secretome findings (Fig. 6). C) Log min-max normalized heatmap of 48-analyte serum Luminex panel performed on terminal serum samples; individual animal replicates are shown rather than condition averages, to convey biological variance associated with progressive moribund state. The in vivo serum Luminex panel (Mouse 48-plex Procarta, cat# EPX480-20834-901) includes analytes not present in the in vitro murine panel (IL-1A, M-CSF, Eotaxin/CCL11, IL-28, IL-18, BAFF/TNFSF13B, MIP1B/CCL4, IL-7); concordance analysis is restricted to the overlapping analyte set.

Supplementary Figure 7

**A**

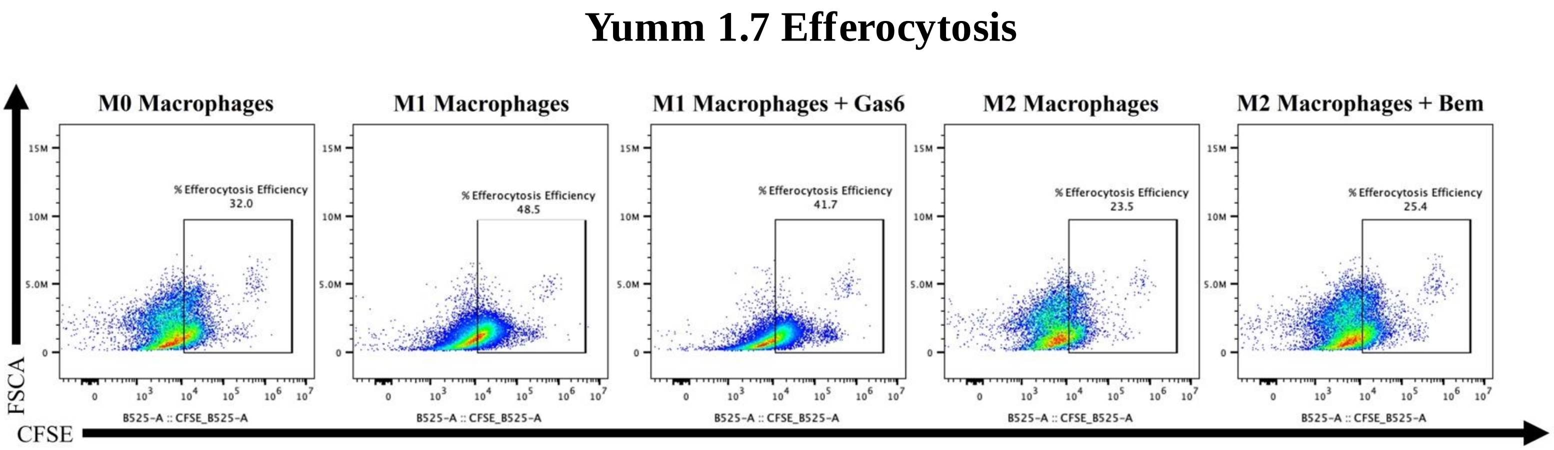


**B**

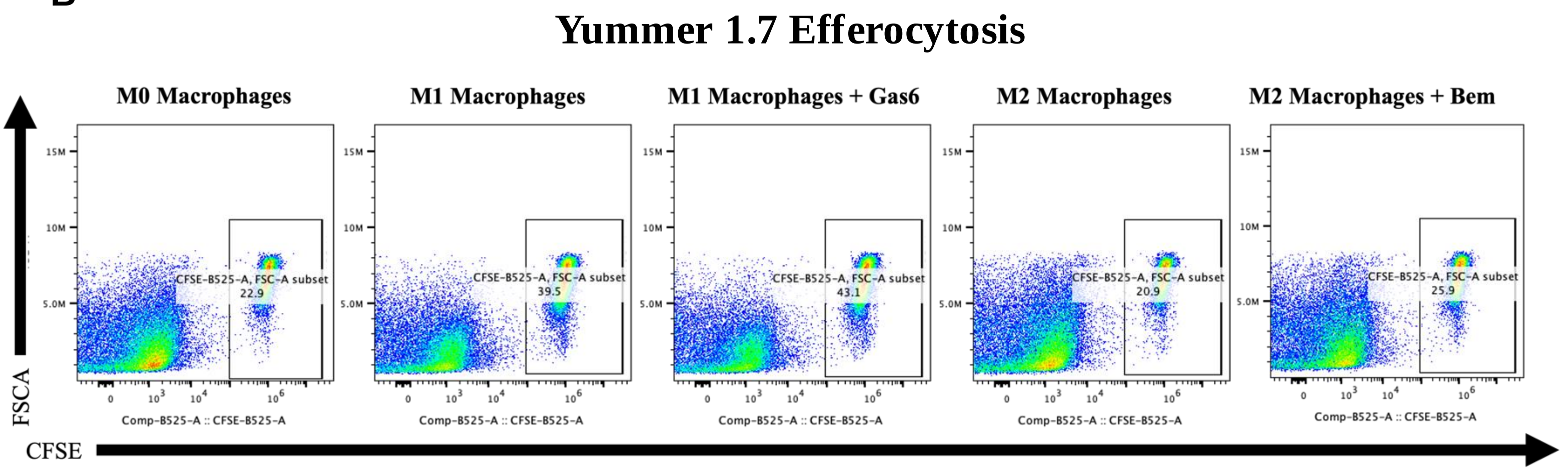

**Supplementary Figure S7. Flow cytometry gating strategy for YUMM1.7 and YUMMER1.7 cell lines.** Representative gating panels demonstrating AXL expression and immune cell identification strategy used throughout in vitro and in vivo experiments.

Supplementary Figure 8

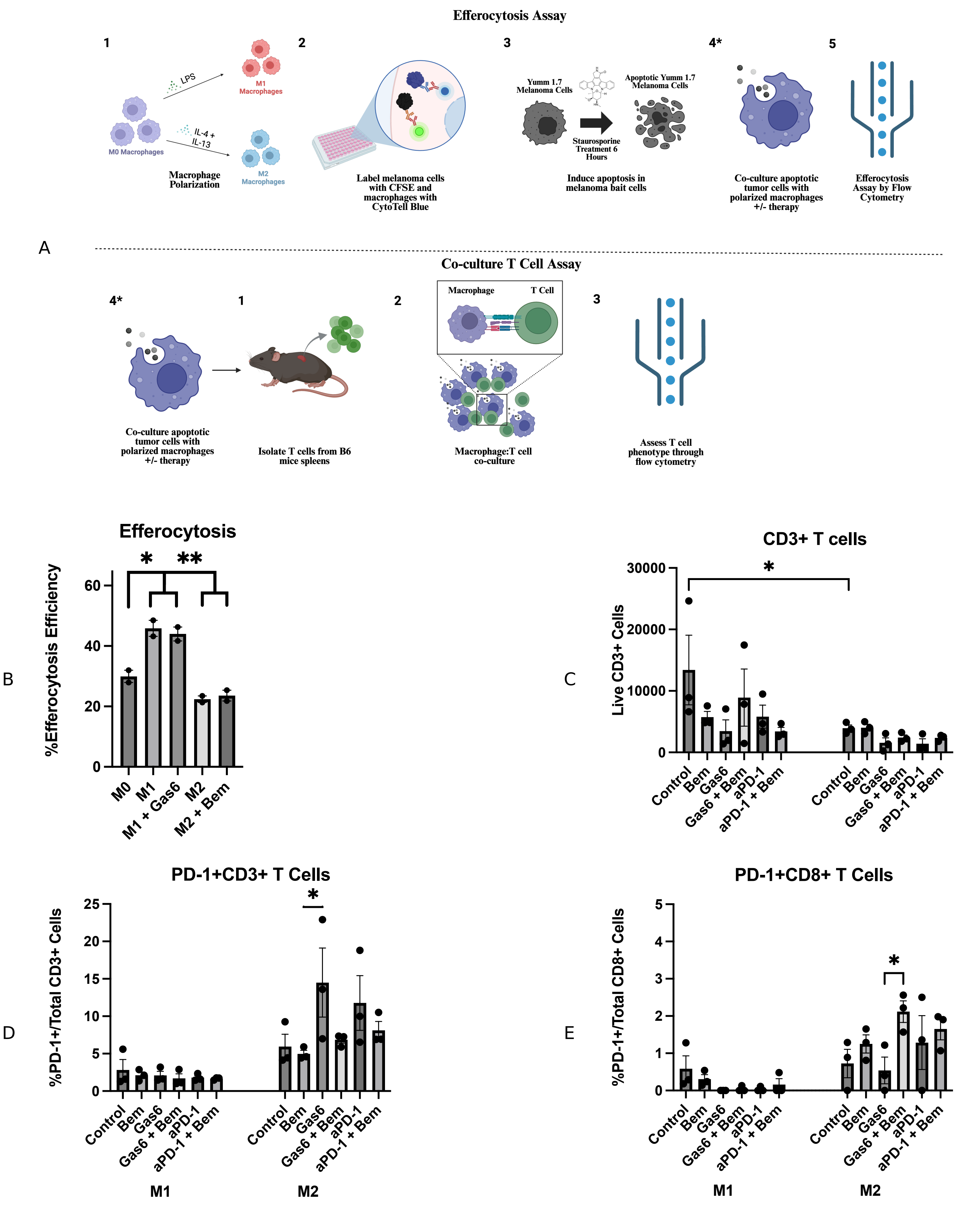

**Supplementary Figure S8. Efferocytosis and T cell co-culture functional assays.** A) Flow cytometry was performed to assess % efferocytosis in a co-culture system through double positivity of CFSE and Cytotel Blue with YUMM1.7 tumor cells. No significant differences in efferocytosis were observed with gas6 stimulation or AXL inhibition in either M1-like or M2-like macrophages, indicating that AXL inhibition does not operate primarily through modulation of phagocytic function. B) Efferocytosis assay repeated with YUMMER1.7 (AXL-low, PD-1 sensitive) tumor cells with similar results. C) CD8+ and CD4+ T cell populations showed no significant differences, excepting decrease in CD8+ T cells with unopposed gas6 exposure, compared to aPD-1+Bem combination in M2-like conditions. D) PD-1+CD3+ and E) PD-1+CD8+ T cell populations under indicated conditions.

Supplementary Figure 9

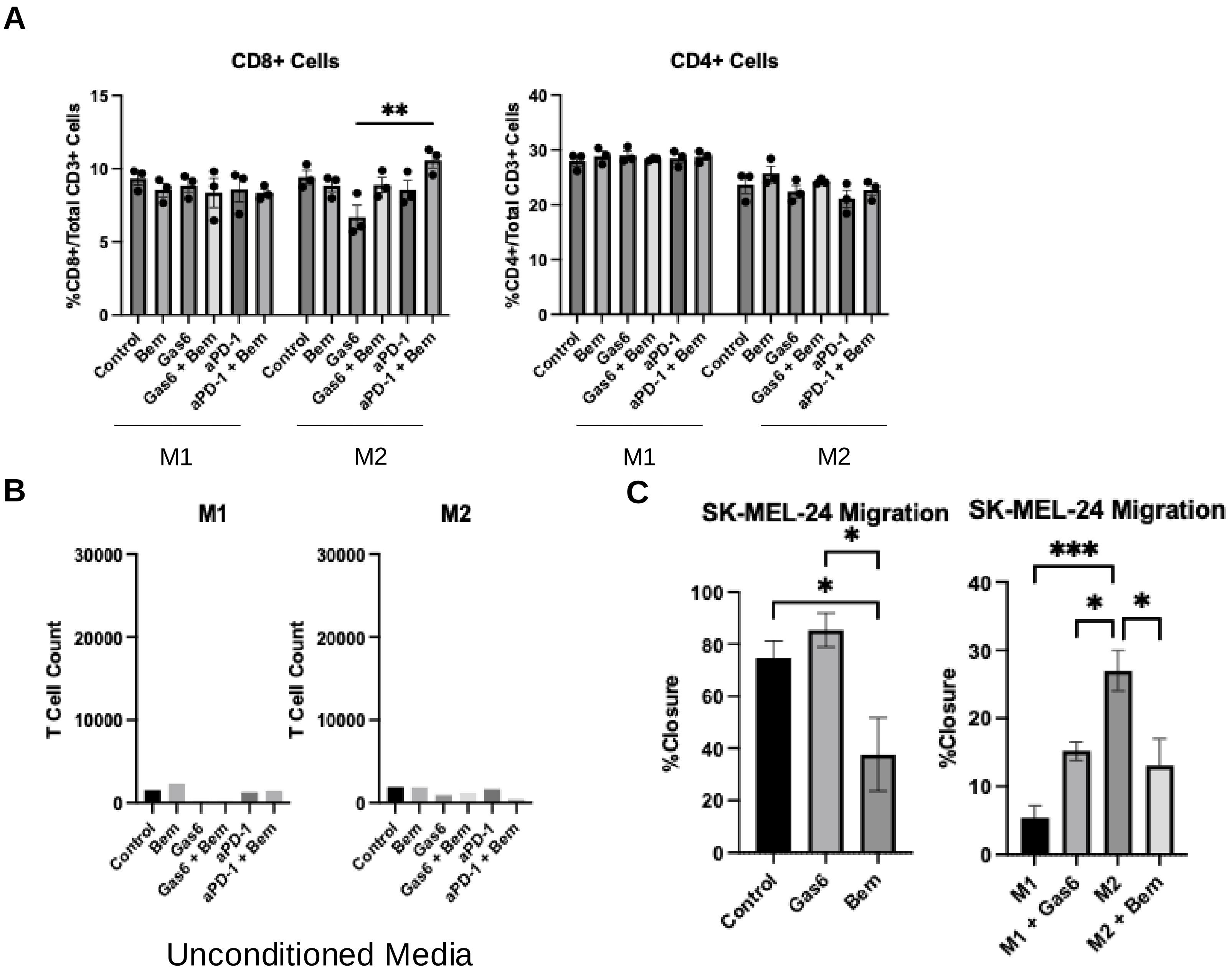

**Supplementary Figure S9.** Macrophage paracrine functions T cells and tumor cells. A) CD8+ and CD4+ T cell populations showed no significant differences, excepting decrease in CD8+ T cells with unopposed gas6 exposure, compared to aPD-1+Bem combination in M2-like conditions. B) Assay was repeated with serum free media, demonstrating no measurable difference in the absence of co-stimulatory factors*.* C) Migration assays were compared with SK-MEL-24 cells treated directly with gas6 or Bem, compared to conditioned media from treated macrophages. Gas6 increased tumor cell migration, while bemcentinib decreased it. M2 secreted factors enhanced tumor cell migration more than M1, with AXL stimulation or inhibition increasing or decreasing migration, respectively.
